# Defined synthetic microbial communities colonize and benefit field-grown sorghum

**DOI:** 10.1101/2023.05.30.542977

**Authors:** Citlali Fonseca-García, Andrew Wilson, Joshua Elmore, Dean Pettinga, Ryan McClure, Jackie Atim, Julie Pedraza, Robert Hutmacher, Robert Egbert, Devin Coleman-Derr

## Abstract

The rhizosphere represents a dynamic and complex interface between plant hosts and the microbial community found in the surrounding soil. While it is recognized that manipulating the rhizosphere has the potential to improve plant fitness and health, engineering the rhizosphere microbiome through inoculation has often proved challenging. This is in large part due to the competitive microbial ecosystem in which the added microbes must survive, and lack of adaptation of these added microbes to the specific metabolic and environmental pressures of the rhizosphere. Here, we constructed an inoculation formula using a defined synthetic community (dSynCom) approach that we hypothesized would improve engraftment efficiency and potentially the relationship with the host plant, *Sorghum bicolor*. The dSynCom was assembled from bacterial isolates that were either: 1) identified to potentially play a role in community cohesion through network analysis, or 2) identified to benefit from host-specific exudate compounds. Growth of the dSynCom was first evaluated *in vitro* on solid media, secondly *in planta* under gnotobiotic laboratory conditions, and finally using sorghum plants grown in the field. We demonstrate that the dSynCom performs best in terms of maintaining diversity when grown in the presence of the plant host in lab conditions, and that many lineages are lost from the community when grown either *in vitro* or in a native field setting. Finally, we demonstrate that the dSynCom is able to promote growth of above- and below-ground plant phenotypes compared to uninoculated controls, both in the lab and when applied to plants grown in the field. These results demonstrate the potential utility of SynComs for supporting crop performance even in the absence of persistence, and the need for a deeper mechanistic understanding of community control of host fitness in agricultural contexts.

## INTRODUCTION

In nature, microbes associated with plants and other eukaryotic organisms self-assemble into communities with remarkable genetic and physiological diversity, often impacting their hosts in complex ways based on forces and factors that we are only beginning to understand (Theis et al., 2016; Compant et al., 2019; Trivedi et al., 2020). Synthetic communities (SynComs), in which strains are assembled together following novel compositional formulas that either mimic or build upon patterns observed in nature, can be used in a number of ways to explore these forces and to understand the additive and synergistic properties that emerge from these communities as they colonize their host. For example, SynComs can serve as testbeds for exploring hypotheses regarding the molecular underpinnings of plant-microbe and microbe-microbe communication (Schäfer et al., 2022), or alternatively to explore the evolution of the microbiome structure and function over time. Additionally, they can also be used to try to augment the colonization ability of individual strains, and to support the development or manipulation of complex phenotypes in their hosts that are challenging to produce with single organisms (Ahkami et al., 2017; Vorholt et al., 2017; Shayanthan et al., 2022).

While interest in use of microbial amendments to support plant performance in agricultural contexts has grown in recent years, development of products capable of robust phenotypic change in field environments has frequently proven challenging. Many such amendments comprised of single isolate strains, where competition from native microbes and environmental perturbation are minimal, have appeared promising in laboratory or greenhouse conditions. However, upon testing in a native field setting, both colonization efficiency and gained benefit are often lost (Deng et al., 2019; Liu et al., 2022; Schmitz et al., 2022). The use of intentionally designed SynComs has been proposed as a promising alternative to single strain inocula (Vorholt et al., 2017) due to the potential for within community positive interactions that bolster colonization and interaction with the host. Currently, additional experimentation is needed to explore plant-associated SynCom performance across a range of environments and the specific biotic and abiotic conditions necessary to foster beneficial impact on their host.

In this study, we aimed to design a defined SynCom (dSynCom) with the ultimate goal of testing engraftment and impact on host phenotype in the bioenergy, feed, and forage crop *Sorghum bicolor*. Sorghum serves as an excellent host for this work as there exists extensive sequencing-based characterization of its microbiome under a range of abiotic conditions and environments (Xu et al., 2018; Muratova et al., 2020; Yukun et al., 2021; Abera et al., 2022; Qi et al., 2022). Using these datasets, and a culture collection of sorghum root-derived bacterial isolates, we selected a total of 57 strains following two complementary but orthogonal guiding principles for inclusion: 1) network-analysis based identification of strains related to abundant and common hub taxa within the sorghum rhizosphere microbiome, and 2) identification of stains capable of using sorghum-specific root exudate compounds as a carbon source. We applied this assembled dSynCom to media-based, gnotobiotic plant-based, and native field-grown plant environments and explored properties of community assembly in each. We observed that while engraftment of the plant root by the dSynCom was undetectable in the field environment, sorghum phenotypes in both growth chamber and field environment were positively influenced by dSynCom treatment. Collectively, these results suggest the promise of SynCom-based strategies for improving agricultural activity and a crucial need for improved understanding of forces controlling engraftment persistence in native environments.

## METHODS

### Network analysis of 16S rRNA data for dSynCom formulation

Strains for the dSynCom formula were identified using two methods. First, we selected strains based on a network analysis of a previously published 16S rRNA dataset (Xu et al., 2018), which originated from an analogous field experiment conducted in 2016 at the same location as the current field experiments (described below). A subset of the previously published 16S rRNA dataset (Xu et al., 2018) that included only the watered rhizosphere samples in the experiment (n=126) was processed as previously described to deliver an ASV occurrence table for downstream analysis that contained approximately 12,000 ASVs (Supplemental Figure 1). Prior to network analysis, the ASV table was filtered to include only ASVs found in at least 50% of samples (n=1762), and then again to include ASVs found only in the top 50th percentile based on average abundance across all samples (n=881). The resulting table was subjected to a Pearson Correlation analysis using CSS normalized count data and the cor function (Bolar, 2019) in R-v4.2.1 (R Core Team, 2022), and ASVs with strong correlations (absolute abundance of greater than 0.8) were maintained in the list of candidate lineages (n=440). A comparison of the taxonomic assignment of the remaining ASVs to an existing sorghum root isolate collection developed in house produced isolate level matches at the genera level for approximately 70% of selected ASVs (n=293). This list was further reduced to a subset of these strains for inclusion in the dSynCom (n=42) by downselection based on availability within our sorghum root isolate collection and abundance (Supplemental Table 1, Supplemental Figure 2).

### Isolation of exudate utilizing microbes from sorghum rhizosphere

Additional strains for the dSynCom formula were selected based on ability to utilize sorghum exudates such as sorgoleone, an allelopathic compound produced at high abundance within the root exudates of sorghum (Yang et al., n.d.; Weston et al., 2013). For this process, field soil was collected from the Kearney Agricultural Research and Extension (KARE) Center in Parlier, California collected at the time when no sorghum was growing and stored at 4°C. MOPS Media Enhanced (MME) medium supplemented with 2 mM single sorghum exudate was utilized for initial enrichment of microbes capable of utilizing sorghum exudates as a sole carbon source. Glass test tubes were used for all cultivation containing any sorghum exudate to minimize adherence of the substrate to the walls of the test tubes. Five mL of this medium was inoculated with 100 mg soil and incubated for 96h. This initial enrichment culture was diluted 1:100 into fresh MME containing 2 mM sorghum exudate supplemented with Wolfes’s vitamins and incubated for 72 h. Following cultivation, 100 µL of the enrichment culture was spread onto MME agar plate containing 2 mM sorghum exudate and incubated for 96 h at 25°C. Well-separated, single colonies were picked and streaked onto R2A agar plate for single colony isolation. Selected isolates were inoculated into MME containing 1 mM sorghum exudate. Isolates that appeared to utilize sorghum exudates were serially passed (1:100 dilution) 6 times into fresh MME containing 1 mM sorghum exudate. These cultures were further spread onto R2A agar plate, picked single colony, inoculated into R2A, and finally pure cultures were stored at −80°C freezer. The 16S rRNA sequences of the isolates were determined via Sanger sequencing, and 15 additional strains were selected from this resource for inclusion in the dSynCom formulation (Supplemental Table 1, Supplemental Figure 2).

### SSM media formulation

To simulate the carbon sources available in the sorghum rhizosphere, a new media recipe was designed based on previously reported metabolomics analysis of the rhizosphere soil of sorghum grown plants (Caddell et al., 2020). The base formula represents a MOPS Media Enhanced (MME) solution, to which a select group of carbon and nitrogen sources were added. The final composition of this media, referred to as Synthetic Soil Media (SMM), is listed in table-format in the Supplementary Materials (Supplementary Table 2).

### Generation of dSynCom and media experiments

To generate dSynCom material for use in media-based experiments, we collected approximately the volume of one inoculating loop (10 uL) of biomass per strain after that strain was grown on plates for 24-96 hours. The amount of time each strain was grown on the plates was based on preliminary experiments designed to determine the amount of time required for each strain to produce sufficient biomass. A total of 57 strains were used to make the dSynCom, 42 strains from network analysis and 15 strains from the isolation of sorghum-exudate utilizing microbes, where some strains were members of the same species. Each strain was plated out on the days described in Supplementary Table 1. Each loopful was then resuspended in 200 uL of sterile 1X Phosphate Buffered Saline (PBS). For each resuspended strain 50 uL was collected and combined into a master mix of all 57 strains. From this mixture 500 uL was plated onto each of the following agar plate types: SSM agar and SSM agar plate with 1:5 diluted sorghum exudates (SSM-Ex). The resulting dSynCom-inoculated plates were allowed to grow at 20 °C for one week before biomass was assessed and collected (Figure 1A). Growth was collected from each plate and resuspended in 2 mL of 1X PBS. This resuspension was then used to make dilutions ranging from 10^-1^ to 10^-2^. With four plate types and two dilutions levels this led to eight dilutions in total. A total of 500 uL of each dilution from a certain plate type (two dilutions per plate type) was then replated onto the same plate type (e.g. all two dilutions from the SSM agar plates were replated onto SSM agar plates). Each of these eight plates were then allowed to grow at 20 °C for one week before growth was collected for 16S rRNA analysis and replated onto the same plate type. This was continued for 7 more passages (for a total of eight passages).

**Figure 1.**
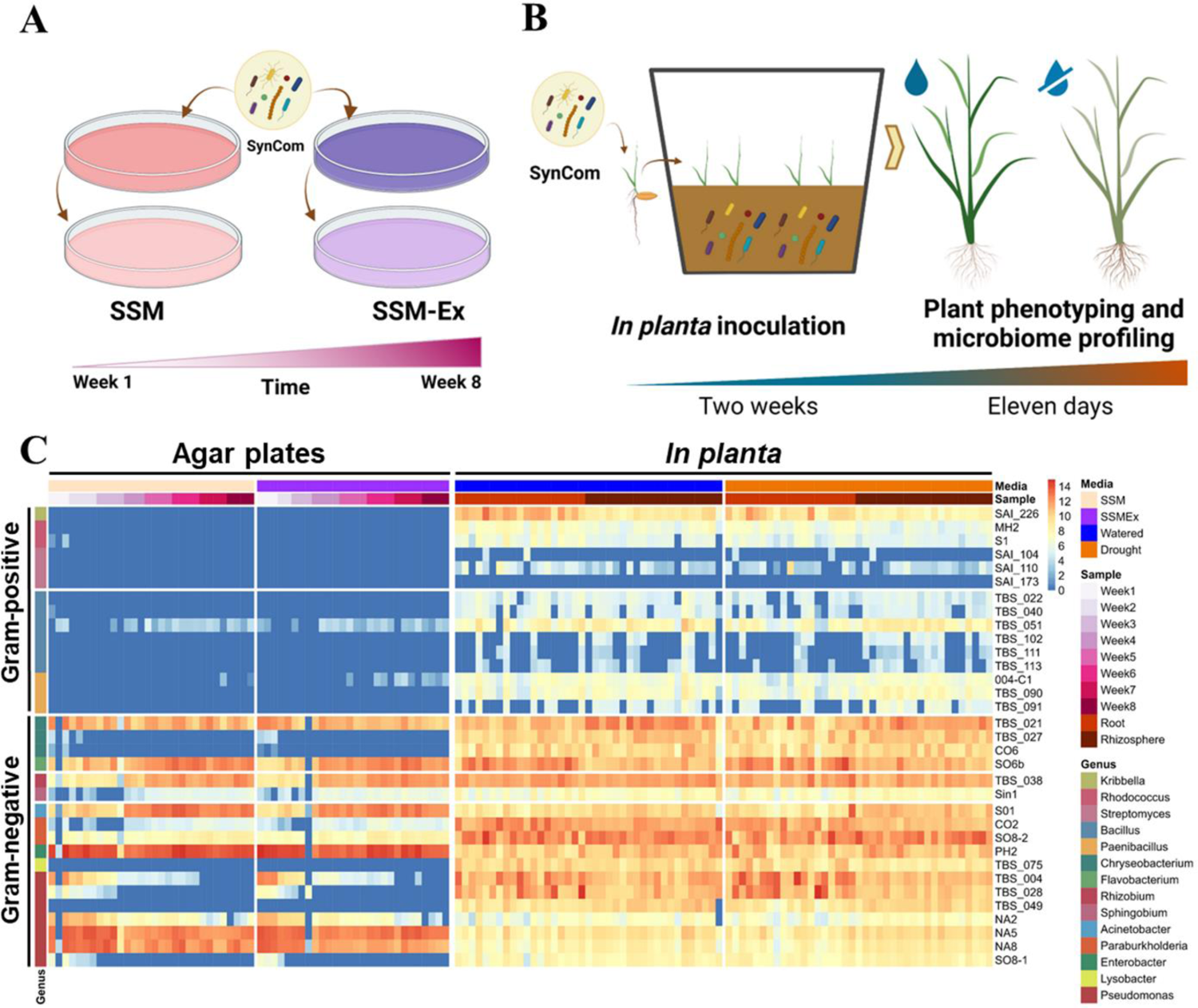
dSynCom colonizes the rhizosphere and roots of sorghum. A) Schematic diagram of the dSynCom stability analysis on agar plates. Dilutions of 10^-1^ and 10^-2^ of the dSynCom were plated on agar plates with Synthetic Soil Media (SSM) or SSM with sorghum exudates (SSM-Ex) and passed over eight weeks. 16S rRNA profiling was performed of at least two replicas across passages. B) Schematic representation of the lab-based dSynCom colonization analysis. Surface disinfected sorghum seeds (cultivar RTx430) were germinated and treated with mock or dSynCom in 5L-capacity microboxes under a sterile environment. Plants were irrigated every three days (26 days) or drought stressed (no irrigation added, 11 days). Plant phenotyping and 16S rRNA profiling were performed at the time of harvest. C) Heat map of the abundance of the dSynCom members identified across treatments *in vitro* and *in planta* experiments. The abundance is presented in log2 scale for n=174 samples for *in vitro* experiments and n=154 samples for the *in planta* experiment.

### Lab-based *in planta* experiments

To explore dSynCom growth *in planta*, sorghum seeds from cultivar RTx430 were sterilized by 10% bleach (sodium hypochlorite) treatment for 15 min, washed five times with sterilized water, and germinated on sterilized wet paper filter at 30 °C without light. To produce enough dSynCom inoculum, each strain was grown on TSA plates (15 g pancreatic digest of casein; 5 g peptic digest of soybean meal; 5 g sodium chloride; 15 g agar; autoclaved once at 120 degrees Celsius for 15 min) at 30 °C according to its grow rates (Supplementary Table 1). We collected approximately six inoculating loopfuls of growth for each strain and suspended in 1.2 mL of sterile 1X PBS. Cell suspensions were pooled and split into 50 mL beakers with 6 mL each. 5L-capacity Microboxes (Combiness, Nevele, Belgium) were used to create a sterilized environment for growth of the sterile seedlings and dSyncom inocula. Microboxes containing 1 kg of calcined clay (Sierra Pacific Supply http://www.sierrapacificturf.com) per box were autoclaved and placed into a laminar hood. A total of 400 mL of autoclaved water with Hoagland’s solution (1.6g/L, Catalog No. H2395-10L, Sigma, San Luis, Misuri) was added into the calcined clay and mixed thoroughly. In each Microbox, we placed four pre-sterilized germinated seeds following a 10 minute incubation with either the dSynCom cell suspension or mock treatment (1X PBS). Each seedling was transplanted 5 cm beneath the soil surface and an additional 1 mL of the dSynCom inoculum or mock treatment buffer was pipetted on top of each seed. Seedlings were then covered with soil. A total of 20 microboxes were used for each experiment: 5 replicas x 2 water treatments x 2 inoculation treatments (mock and dSynCom). Plants were irrigated with 3 mL of sterile Hoagland’s solution per plant three times a week (Figure 1B). After two weeks the lids were removed, and the drought treatment was applied for 11 days. Growth conditions were 33°C/28°C, 16h:8h Light/Dark, 60% humidity for 26 days before harvesting. At collection, shoot phenotyping was performed and roots were used for microbiome profiling.

### Field Experiment

To explore dSynCom growth in field grown plants, a field experiment was conducted at the Kearney Agricultural Research and Extension (KARE) Center in Parlier, California (36.59, - 119.51). Sorghum cultivar RTx642 seeds were planted (June 9, 2022) within a block design and two different types of irrigation (drought stressed and watered plants) with three replicate blocks per treatment (Supplemental Figure 3). Prior to planting, all sorghum seeds were inoculated with either the dSynCom or mock treatment. Sets of 200 seeds were incubated into 50 mL conical tubes with 15 mL of dSynCom or mock (PBS 1X buffer) for 45 min at room temperature and then dried for 3 hours in a laminar hood. dSynCom preparation was carried out under the same conditions as described for the Microbox experiments above, with one modification: we added 0.01% tween 20 to the 1X PBS as an adherent. Each block had four rows treated with mock (R2-R5) and four rows treated with the dSynCom (R6-R9). Unfortunately, one block of each plot presented a low rate of germination, leaving only four blocks in total for the experiment. Five <u>w</u>eeks <u>a</u>fter <u>p</u>lanting (<u>wap</u>), a second inoculation (dSynCom or mock) was applied to 10 selected plants per row from each block, and these plants were labeled with numbered stakes. Plants of a similar size and with only one to two tillers were selected; inoculation consisted of 1 mL of dSynCom diluted into 200 mL of tap water. At 9 wap, a drought treatment was imposed on one half of the blocks, consisting of a complete lack of irrigation until rewetting following the microbiome sample collection (Supplemental Figure 3). Watered blocks, which account for the remaining half of the blocks, received irrigation for the entire growing season beginning in the third wap; no water was applied in the first three weeks to allow seedling establishment. At the end of the drought treatment (9 wap), a total of 160 plants were sampled for microbiome profiling and plant phenotyping. These consisted of 5 plants inoculated from each treatment group (2 inoculation types x 2 inoculation times x 2 water treatments x 2 replicate rows x 2 replicate blocks). At 16 wap, an additional 10 plants from each treatment group (n=320) were bagged to protect the seed development, and later harvested for shoot phenotyping and yield quantification at 25 wap.

The amount of water applied to watered plants (3-25 wap) and drought blocks (3-5 and 9-25 wap) was 80% of calculated evapotranspiration each week. Daily potential evapotranspiration (ETo) was determined from an on-site weather station located approximately 1200 m from the field site. A locally-derived crop coefficient was matched to the crop growth stage and multiplied by the ETo values to determine a calculated daily estimated crop evapotranspiration (ETc) for a non-stressed grain sorghum crop. A drip system was utilized to apply all irrigation water during the growing season, consisting of drip lines placed on the surface of each furrow (0.76 m row spacing), with 0.3 m emitter spacing and 2 L/h emitter output. The drip system was only operated for part of one day out of each 7-day period, with treatments being irrigated for that period receiving an amount equivalent to 100 percent of the estimated ETc calculated for the prior 7-day period. Surface drip lines were used for irrigation to provide accurate water application amounts and a high level of water application uniformity. The irrigation treatment designed to be a non-water-stressed block was irrigated at 7-day intervals between the first within-season irrigation (June 9) and the final irrigation (December 1). Measured rainfall during the June 9 to December 1 period was less than 7.11 mm.

### Sample Collection

Plants of lab-based *in planta* experiments were harvested at the end of the drought treatment. Plants were carefully placed and separated in a disinfected tray with new aluminum foil. Rhizosphere samples were collected using 8 ml of epiphyte removal buffer (0.75% KH2PO4, 0.95% K2HPO4, 1% Triton X-100 in ddH2O; filter sterilized at 0.2 μM) with manual agitation and centrifuged to pellet the resulting rhizosphere soil at 3700 rpm for 10 min. Shoots and roots were separated with sterile scissors and roots were stored together with rhizosphere at −80 °C until DNA extraction.

Plant samples from the field trial were collected by manually extracting whole plants with root systems using a shovel to a depth of approximately 30 cm. Rhizosphere and root samples were collected (soil tightly adhering to the sorghum root surface) by pooling multiple roots from each individual plant, for a total of 20 rhizosphere/root samples per treatment type at 9 wap. Roots were frozen on dry ice and then transported to the laboratory (all samples were collected between 9am and 1pm during a single day). In the lab, roots were vortexed in 30 ml of cold epiphyte removal buffer for 5 min and centrifuged to pellet the resulting rhizosphere soil at 3700 rpm for 10 min after removal of the root tissue. Root endosphere samples were obtained by washing the vortexed roots three times in fresh sterile water. To aid in the DNA extraction process, roots samples were first ground in a mortar and pestle with liquid nitrogen and then stored at −80 °C.

### Plant Phenotyping

Plant phenotypic data was recorded at 9 and 25 wap of the field trial. At 9 wap, plant height was recorded two days prior to the microbiome sample collection day. At sample collection, shoot tissue was separated from harvested root tissue with 70% ethanol disinfected pruning shears and shoot fresh weight biomass was measured. The shoot tissue was subsequently dried in paper bags at 50 °C for 14 days and weighed again for dry weight. Additional root phenotyping of the root ball was performed after root and rhizosphere sampling using the RhizoVision Crown system following the protocols established in Seethepalli et al., (2020). As described above, at 16 wap 10 plants per row were bagged for 10 weeks to avoid birds and animal predation. At 25 wap, the plant height of bagged plants was recorded and each bagged plant was then harvested. Shoots were separated from roots and panicles and weighed in a balance, dried in paper bags at 50 °C for 14 days, and weighed again. To quantify yield production, the seeds of each panicle were threshed a week after harvest and weighed. A thousand seeds of each panicle were then counted and weighed to establish 1000 seed weight.

Statistical analyses of the shoot phenotyping data were performed in R-v4.2.1 (R Core Team, 2022). Significance was tested by ANOVA one-way, Brown-Forsythe and Welch ANOVA one-way, and Kruskal Wallis according to their distribution and variance homoscedasticity using the aov, oneway.test, and kruskal.test functions in the R stats package (Bolar, 2019). Distribution and variance homoscedasticity were tested using the shapiro.test and bartlett.test functions also in the R stats package (Bolar, 2019). Post-hoc pairwise comparisons tests were conducted using HSD.test function in the R package Agricolae (de Mendiburu, 2021), pairwise.t.test function in the R stats package (Bolar, 2019), and kwAllPairsDunnTest function in the R PMCMRplus package (Pohlert, 2022). Root phenotyping data was analyzed using pca function in R PCAtools package (Blighe K, 2023) and biplot function in R stats package (Pohlert, 2022), and treatment significance was determined by PERMANOVA test with 999 permutations using adonis function in R vegan package (Dixon, 2003).

### DNA Extraction and Library preparation

DNA extraction of root and rhizosphere samples was performed using Qiagen DNeasy Powersoil DNA extraction Kit (Catalog No. 12888-100; QIAGEN, Venlo, Netherlands) with 150 mg of root and 200 mg of rhizosphere as starting material in the provided Powersoil collection vials and then following the manufacturer’s directions. DNA concentrations were measured with a Qubit 3.0 Fluorometer and samples were diluted to the same template amounts. Samples were amplified using a dual-indexed 16s rRNA Illumina iTags primer set specific to the V3-V4 region 341F (5’-CCTACGGGNBGCASCAG-3’) and 785R (5’-GACTACNVGGGTATCTAATCC-3’) using Invitrogen Platinum Hot Start PCR Master Mix (2X), (Catalog No. 13000014, Waltham, Massachusetts) and 10 ng of template per reaction. PNAs (Lundberg et al., 2012) designed to target host derived amplicons from chloroplast and mitochondria 16S rRNA sequences were added to root reactions (0.75 µM of each, PNABIO, Thousand Oaks, CA). PCR reactions were performed in triplicate in three individual thermocyclers to avoid possible thermocycler bias with the following conditions: initial cycle of 3 min at 98°C, then 30 cycles of 45 s at 98°C, 10 s at 78°C, 1 min at 55°C, and 1.5 min at 72°C, followed by a final cycle of 10 min at 72°C. Triplicates were then pooled and DNA concentration for each sample was quantified using Qubit 3 Fluorometer. Pools of amplicons were made using 100 ng for each PCR product. 600 ng of pool amplicons in 100 µl of volume were cleaned up with 1.0X volume of AMPureXP (Beckman-Coulter, West Sacramento, CA) beads according to the manufacturer’s directions. An aliquot of the pooled amplicons was diluted to 10 nM in 30 µl total volume before submitting to the QB3 Vincent J. Coates Genomics Sequencing Laboratory facility at the University of California, Berkeley for sequencing using Illumina Miseq 300 bp pair-end with v3 chemistry.

### Amplicon sequence processing and analysis

16S amplicon sequencing reads were demultiplexed in QIIME2 (Bolyen et al., 2019) and then passed to DADA2 (Callahan et al., 2016) to generate Amplicon Sequence Variants (ASVs), with taxonomies assigned using the December 2019 version of Silva 16S rRNA gene 138 database. All subsequent 16S statistical analyses were performed in R-v4.2.1 (R Core Team, 2022). In both the media based and *in planta* experiments, we prefiltered out ASVs that were not alignable via BLAST and phylogeny tree to at least one of the 57 16S rRNA sequences of the dSynCom members. Before filtering, we observed 175 ASVs in both data sets with more than 10 reads per sample, of which 102 ASVs were detected as plant contaminants and 21 ASVs as agar plate contaminants. The detected contamination corresponded only to 1.27% of the total amount of reads and the remaining 98.73% of reads were assigned to the ASVs of the dSynCom members. In the field experiment, we identified the members of the dSynCom using again BLAST and phylogeny approaches but now with the complete set of ASVs. To account for differences in sequencing read depth across samples in all the experiments, samples were normalized by dividing the reads per ASV in a sample by the sum of usable reads in that sample, resulting in a table of relative abundance frequencies, which were used for analyses. Alpha diversity was determined with the estimate_richness function in the R package phyloseq (McMurdie and Holmes, 2013), and significance was tested by ANOVA using the aov function in the R stats package (Bolar, 2019). Beta diversity (PCoA and CAP) was performed using the ordinate function in the R package phyloseq (McMurdie and Holmes, 2013). Treatment separation was determined by pairwise PERMANOVA with 999 permutations using the adonis and calc_pairwise_permanovas functions in the R packages vegan (Dixon, 2003) and mctoolsr (Leff, 2016). Additionally, the permutest function was used for constrained ordination (CAP) analysis with 999 permutations also in the R package vegan (Dixon, 2003). Tukey-HSD tests were used for the HSD.test function in the R package Agricolae (de Mendiburu, 2021). The bacterial ASV heatmaps were generated using the R package pheatmap (Kolde, 2019). Differentially abundant ASVs were conducted using R package edgeR (Robinson et al., 2010), the ASVs with a log2FoldChange > 1.5, *P* < 0.05 were considered differentially significant.

## RESULTS

In order to develop an inoculation for the rhizosphere environment that would allow us to test permutations of membership and compositional abundance, we chose to formulate a synthetic community of bacterial isolates using a bottom-up approach. To identify the combination of strains that would be well-suited to collectively support colonization and establishment in and on the root, we used a combination of two distinct strain-selection strategies. First, we aimed to identify bacterial lineages that we hypothesized would be connected to community composition integrity by applying a network-based analysis of an existing 16S rRNA microbiome dataset from the sorghum rhizosphere environment. This analysis was performed using a Pearson correlation on 16S rRNA data for a total of 126 rhizosphere samples collected from the KARE field station in 2016 (Xu et al., 2018). Using this approach and additional selection parameters based on ASV abundance and occupancy across the dataset (Supplemental Figure 1), we identified a total of 293 ASVs belonging to 16 genera. A final selection based on availability within our sorghum root isolate collection and abundance was used to narrow the list of target strains to 42 bacterial isolates (Supplemental Table 1). As a second strategy for engineering a robust community capable of colonizing the sorghum rhizosphere, we also aimed to include strains that could effectively use sorghum-produced exudates. To achieve this, we performed isolations of bacterial strains from KARE field sorghum soil that could utilize sorghum exudates as a sole carbon source. These isolations yielded 15 individual strains, belonging to 10 additional genera; we included each of these strains (Supplemental Table 1) in the final dSynCom formulation (Supplemental Figure 2).

### *In Vitro* Experiments

To test the ability of this assembled formulation of strains to grow in co-culture, we first assembled the community and applied it to solid media comprised of synthetic compounds intended to mimic the composition of sorghum root exudates. Each of the 57 individual strains were first grown on SSM and SSM-Ex plates and then mixed using approximately equal volume of cells per strain as input. After incubating, samples were collected weekly to assess community composition and diversity through 16S rRNA (V3-V4 region) sequencing (Figure 1A). The 16S rRNA ASVs identified as corresponding to each dSynCom member across this dataset were grouped according to their similarity in the V3-V4 region as is listed on the Supplemental Table 1. The resulting data demonstrate that approximately 55% of the dSynCom strains were no longer detectable on the media after the second week of growth (Figure 1C). In particular, in this plate-based environment, the majority of strains selected using the network analysis and isolated from the sorghum roots did not persist. Additionally, inclusion of sorghum root exudates in the media collected from 5- to 10-week old plants did not appear to improve survival rates of these root-derived strains, nor did it seem to impact community dynamics over the course of the experiment. Interestingly, the majority of strains selected based on their ability to utilize sorghum exudates as a sole carbon source remained detectable for the duration of the eight-week experiment. These results demonstrate that the intended overall community structure for the dSynCom is not well supported by *in vitro* growth on solid media.

### *In planta* Experiments

We hypothesized that many of the strains that did not persist in the media-based experiments would grow better in the native habitat from which they were isolated, namely the sorghum rhizosphere. To test this hypothesis, we developed a lab-based *in planta* experiment using gnotobiotic containers (microboxes) filled with calcined clay and with either addition of dSynCom or mock inoculation treatment applied to growing seedlings. The dSynCom inocula was prepared using similar steps as described for the *in vitro* experiments and applied to germinated sorghum seeds that had been previously surface disinfected (Figure 1B). Following a period in which all plants were irrigated, one treatment group was subjected to a period of drought stress prior to microbiome analysis and plant phenotype collection. As anticipated, the majority of the dSynCom strains (56 out of 57) were detectable in both root and rhizosphere samples of dSynCom treated sorghum under both watering treatments. Only *Brevibacillus agri* TBS_043 was not detected in any sample (Figure 1C). While sorghum-exudate solubilizers dominated the community composition in media-based growth, these data showed a much more even distribution of abundance across the dSynCom formulation is possible when the plant host is added to the environment.

Apart from dSynCom treatment, other experimental factors appeared less influential in determining community composition. Likely due to the short duration of the drought stress in this experiment, watering treatment did not greatly impact the composition or conformation of the community in terms of alpha and beta diversities (Supplemental Figure 4, Supplemental Table 3). In addition, the plant compartment (rhizosphere or root) lead to only slight differences in composition, with impacts primarily in a select few members of the dSynCom, such as *Kribella shirazensis* SAI_225 and *Flavobacterium anhuiense* SO6b that show higher abundance in roots. Overall, we observed that Gram-negative member strains of the dSynCom remained better colonizers than Gram-positive members, independent of the method used to select them for the community (network analysis or isolated on sorghum exudates containing media). Interestingly, we noted that under watered conditions, application of the dSynCom positively affected plant phenotype compared to the mock treatment; both shoot fresh weight and dry weight increased in dSynCom inoculated plants compared to mock inoculated controls in the watered treatment (Figure 2). Collectively, these results indicate that the dSynCom grows more uniformly in the presence of the plant root and has the ability to improve plant biomass, showing the need for the root environment for a stable assembly which impacts the plant fitness simultaneously.

**Figure 2.**
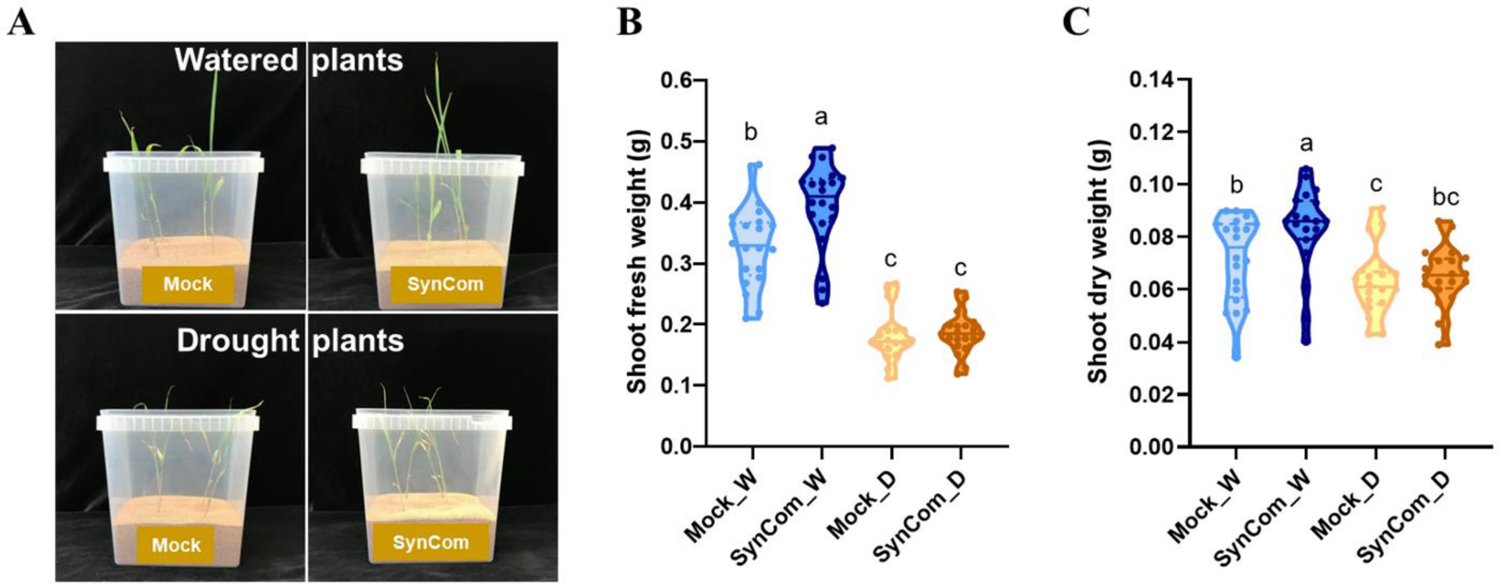
dSynCom increases the shoot biomass of watered sorghum plants in lab-based *in planta* experiments. A) Representative pictures of watered or drought stressed sorghum plants inoculated with mock or dSynCom. B) Shoot fresh weight. C) Shoot dry weight. W=normal irrigation (blue colors); D=drought stress (brown colors). Letters above represent statistical differences among treatments by Brown-Forsythe and Welch ANOVA one-way, Holm-Šídák post hoc test *P* < 0.05 of 18-20 plants per treatment (78 plants in total).

To explore whether the inclusion of sorghum-exudate utilizing strains impacted colonization efficiency of the other dSynCom members, we repeated this *in planta* experiment once more using a modified formula in which the sorghum-exudate solubilizers were removed from the community. Interestingly, we noted that many other community members showed increased representation in the resulting 16S rRNA datasets (Figure 3), including the strains *Streptomyces ambofaciens* SAI_104, *S. albogriseolus* SAI_173/190, *Paenibacillus lautus* TBS_091, and *Pseudomonas plecoglossicida* TBS_049 (Supplemental Figure 5). We also noted that the previously observed positive influence on plant phenotype was conserved even in the absence of the sorghum-exudate utilizers; this was true not only under watered conditions, but also appeared during drought conditions under the new formulation (Figure 4). Collectively, these results show that the inclusion of sorghum-exudate solubilizers impacts the composition of the dSynCom community in sorghum roots and isn’t necessary for the observed impact on plant health.

**Figure 3.**
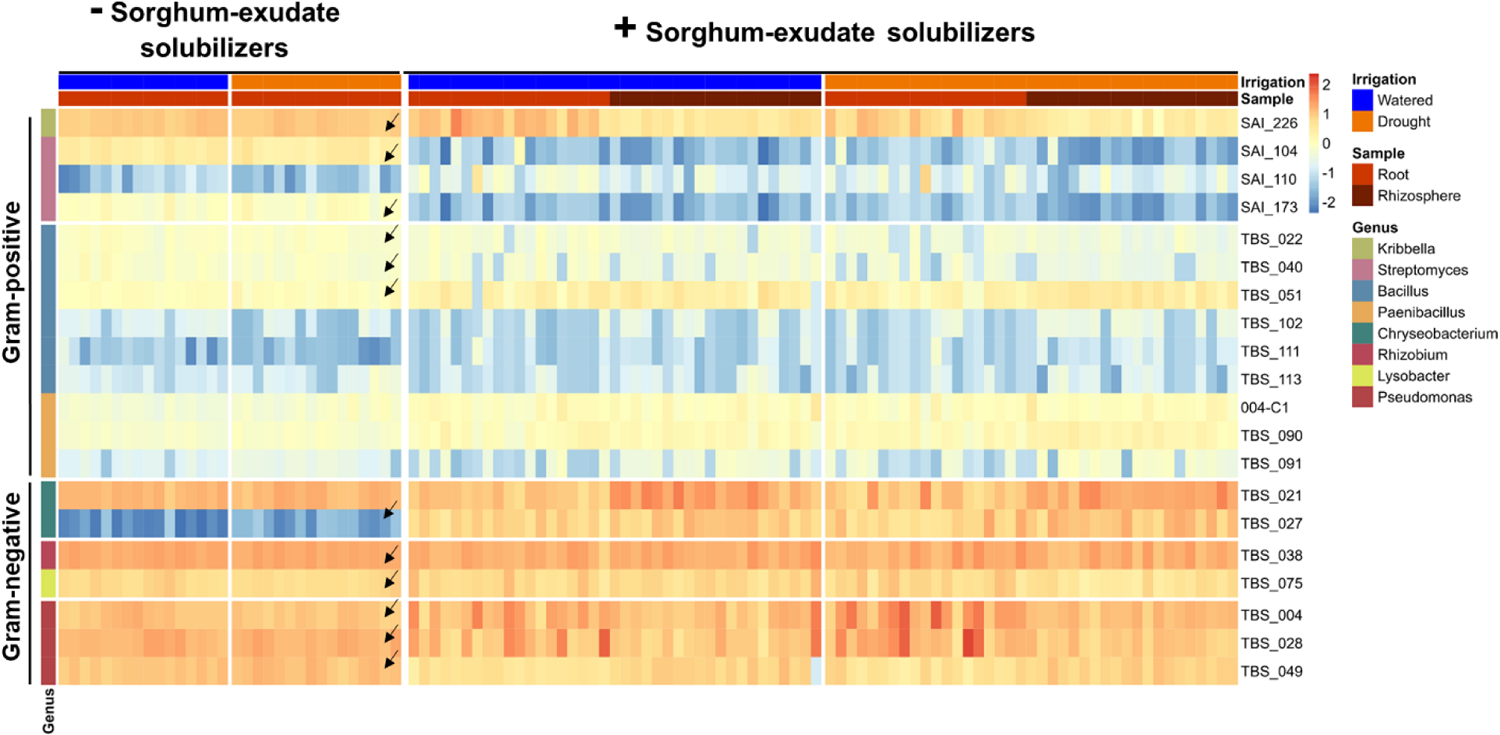
Sorghum-exudate solubilizers modulate the dSynCom conformation. Heat map of the abundance of the dSynCom members with or without sorghum-exudate hydrolyzers identified in lab-based *in planta* experiments. Abundance is presented in log2 scale of n=218 samples. Arrows highlight the dSynCom members potentially modulated by the sorghum-exudate solubilizers after a Brown-Forsythe and Welch ANOVA one-way and Holm-Šídák post hoc test *P* < 0.05. Sample types (roots in red and rhizosphere in dark red) and water treatments (watered in blue and drought in brown) are shown at the top of the heat map. Genera and members present in the dSynCom are shown at the left and right sides, respectively.

**Figure 4.**
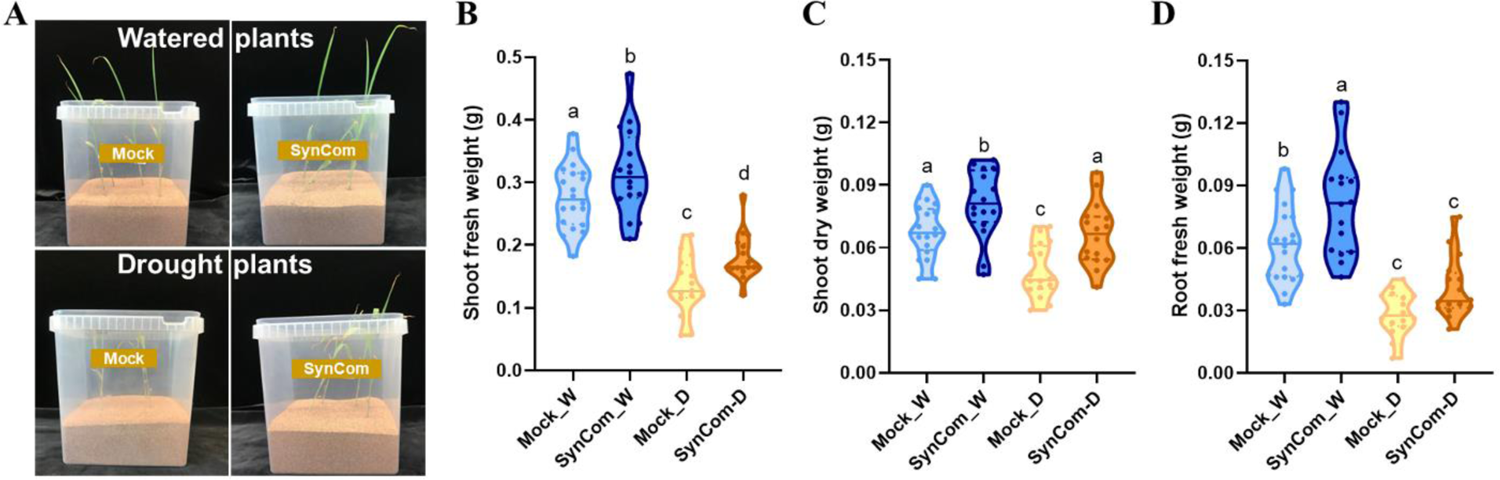
dSynCom, without sorghum-exudate solubilizers, increases the overall shoot biomass of sorghum plants in lab-based *in planta* experiments. A) Representative pictures of watered or drought stressed sorghum plants treated with mock or dSynCom. B) Shoot fresh weight. C) Shoot dry weight. D) Root fresh weight. W=normal irrigation (blue colors); D=Drought stress (brown colors). Letters above represent statistical differences among treatments by ANOVA one way test and Tukey post hoc test with *P* ≤ 0.05 of 16 plants per treatment (n=64 plants in total).

### Field Experiments

The colonization of our dSynCom under controlled and presterilized conditions suggests that this community can effectively assemble on the sorghum root in the absence of competition from other environmental microbes. To explore how the dSynCom performs in a native context, we conducted experiments in the field through inoculation of the dSynCom onto seeds and young growing seedlings. In this experiment, sorghum seeds were first subjected to either a preincubation with the dSynCom treatment or a mock inoculation prior to sowing into the field at the Kearney Agricultural Research and Extension Center. A second round of inoculation (dSynCom or mock) was performed on a select group of 4-week old seedlings in each plot to explore the impact of alternative methods of SynCom deployment. Subsequently, half of the plants in all inoculation treatment types were subjected to a period of drought stress, during which no water was applied for four consecutive weeks (5 - 9 wap). The other half of the plants remained regularly irrigated throughout the experiment (Figure 5A). At the end of the drought treatment (9 wap), we collected root tissue and biomass phenotypes from all treatment groups and performed 16S rRNA (V3-V4) community profiling of root and rhizosphere using Illumina MiSeq sequencing (9 wap). In accordance with previous studies on the sorghum microbiome (Xu et al., 2018), microbial diversity significantly differed between roots and rhizosphere, showing a lower Shannon diversity in roots, and also showed differences according to watering treatment with overall reduced diversity in droughted samples compared with watered treatments (Supplemental Figure 6A). It was also noted that the dSynCom inoculation significantly impacted alpha diversity of watered root samples (under seed inoculation), increasing the Shannon diversity by approximately 2%, from 5 to 5.15 in mock- and dSynCom-treated plants, respectively (no effect was observed in the rest of the treatments).

**Figure 5.**
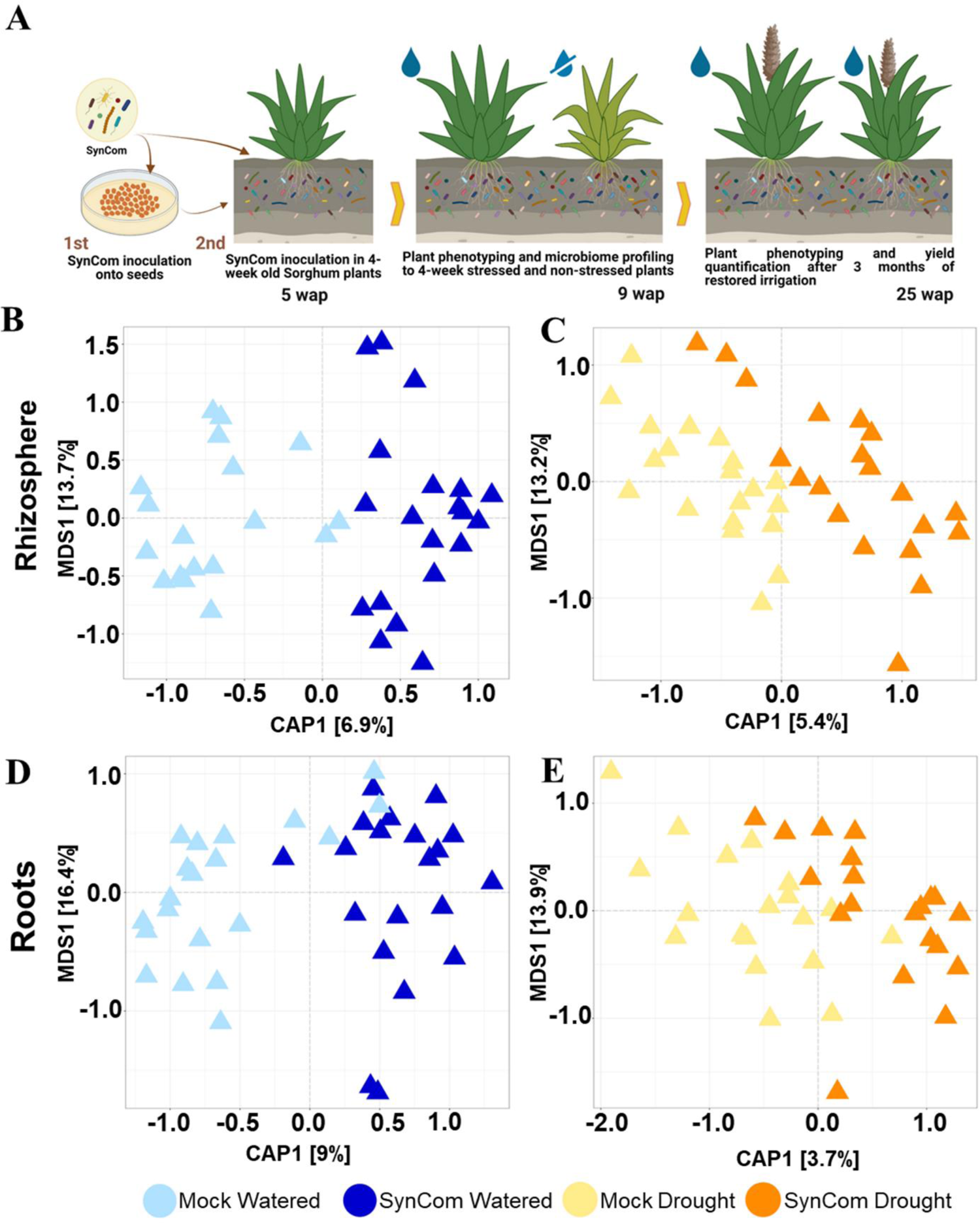
dSynCom impacts the beta diversity of the sorghum root microbiome in the field. A) Schematic diagram of the field trial design. Sorghum plants cultivar BTx642 were treated once (seeds) or twice (seeds and 4-week old plants) with mock or dSynCom. Two irrigation treatments were applied, a drought stress, no irrigation, for four weeks after the second inoculation (5 wap - 9 wap), or irrigation periodically. Microbiome profiling was analyzed at the end of drought stress (9 wap) and the plant phenotyping was recorded also at 9 wap and after three months of the restored irrigation at 25 wap. B) Beta diversity analysis of the sorghum microbiome of the field trial. Constrained Analysis of Principal Coordinates (CAP) ordination plots of the microbial community in the sorghum (B-C) rhizosphere and (D-E) roots samples under (B,D) normal irrigation (blue colors) and (C,E) drought stress (brown colors) inoculated twice with mock or dSynCom.

In order to evaluate the effect of dSynCom treatment on the structure of the microbial community, we analyzed the beta diversity through unconstrained Principal Coordinates Analysis (PCoA) using the Bray–Curtis dissimilarity. Four clusters were grouped across the two axes (Supplemental Figure 6B), where the primary axis clustered samples principally by water treatment (explaining 32.8% of variation), and the second axis by sample type (explaining 10.6% of total variation), in agreement with previous sorghum microbiome studies (Xu et al., 2018). A permutational multivariate analysis of variance (PERMANOVA) analysis was performed using the Bray–Curtis distance for sample type, water treatment, dSynCom/mock treatment and inoculation time points. These analyses indicate that the microbial community was significantly influenced by all four experimental factors (Supplemental Table 4), but mostly by the sample type and water treatment. To analyze the effect of the dSynCom inoculation on community composition in more detail, we performed Constrained Analysis of Principal Coordinates (CAP) using Bray Curtis dissimilarity and Permutation tests of multivariate homogeneity of group dispersions (permutest) for each sample type, water treatment, and dSynCom inoculation time points. Collectively, these analyses indicate that the dSynCom inoculation had a significant but small impact (10.2-6.9% for watered and 6.5-3.7% for drought) on the plant microbiome across all treatments (Figure 5B, Supplemental Figure 7, and Supplemental Tables 4-5). Collectively, these results confirm that sorghum microbiome manifests strong niche differentiation and suggest that the dSynCom impacts the microbial composition.

To explore the impact of each experimental factor on specific taxonomic fractions of the sorghum microbiome, we analyzed the data at the phylum-level and genus-level using relative abundance plots. In agreement with previous studies of root microbiome response to drought stress (Xu et al., 2018), we observed an enrichment of Gram-positive bacteria in drought-stressed samples together with a depletion of Gram-negative bacteria. More specifically, the phylum Actinobacteria and the genus *Streptomyces* were strongly abundant in both roots and rhizosphere of drought stressed samples (Supplemental Figures 8-9), while the phyla Proteobacteria and Bacteroidetes were less abundant in both sample types. Since we had observed a relatively small impact of inoculation timing (seeds versus seedlings) on beta diversity, the remaining microbial composition analyzes were conducted on combined data treating the two inoculation type points as additional replicates.

Interestingly, these analyses also reveal that dSynCom inoculation did not appear to lead to significant increases in the abundances of dSynCom community members within the root or rhizosphere (Figure 6). Given that all dSynCom members were originally isolated from soil and roots obtained from this field environment, it was expected that even mock inoculated samples would show evidence of presence of dSynCom ASVs. However, no consistent, significant enrichment of dSynCom ASV abundance was observed in either root or rhizosphere across our experimental design following dSynCom inoculation compared to the mock control. In light of this, it is all the more surprising that we did observe significant impacts of dSynCom treatment on many distinct taxonomic groups (Figure 7). At the phylum level, a much greater fraction of Firmicute ASVs showed significant decreases in dSynCom treatment compared to the mock control (Figure 7A, Figure 8A). By contrast, ASVs belonging to the phylum Bacteroidetes were more likely to have increased abundance following dSynCom treatment (Figure 7B, Figure 8B). For Actinobacterial ASVs, which tend to be enriched during drought stress, a greater percentage showed gains in abundance following dSynCom treatment under watered conditions, but the opposite pattern was observed under drought conditions (Figure 7C, Figure 8C). At finer taxonomic resolutions, each of these phylum level changes appeared driven by enrichment or depletion of a small and select number of target families (*Bacillaceae* within the Firmicutes, *Chitinophagaceae* within the Bacteroidetes, and *Streptomycetaceae* within the Actinobacteria) (Figure 8D-F). Collectively, these results demonstrate that impact of dSynCom inoculation on root microbiome structure lies largely in rewiring existing community abundance patterns rather than permanent engraftment of dSynCom members.

**Figure 6.**
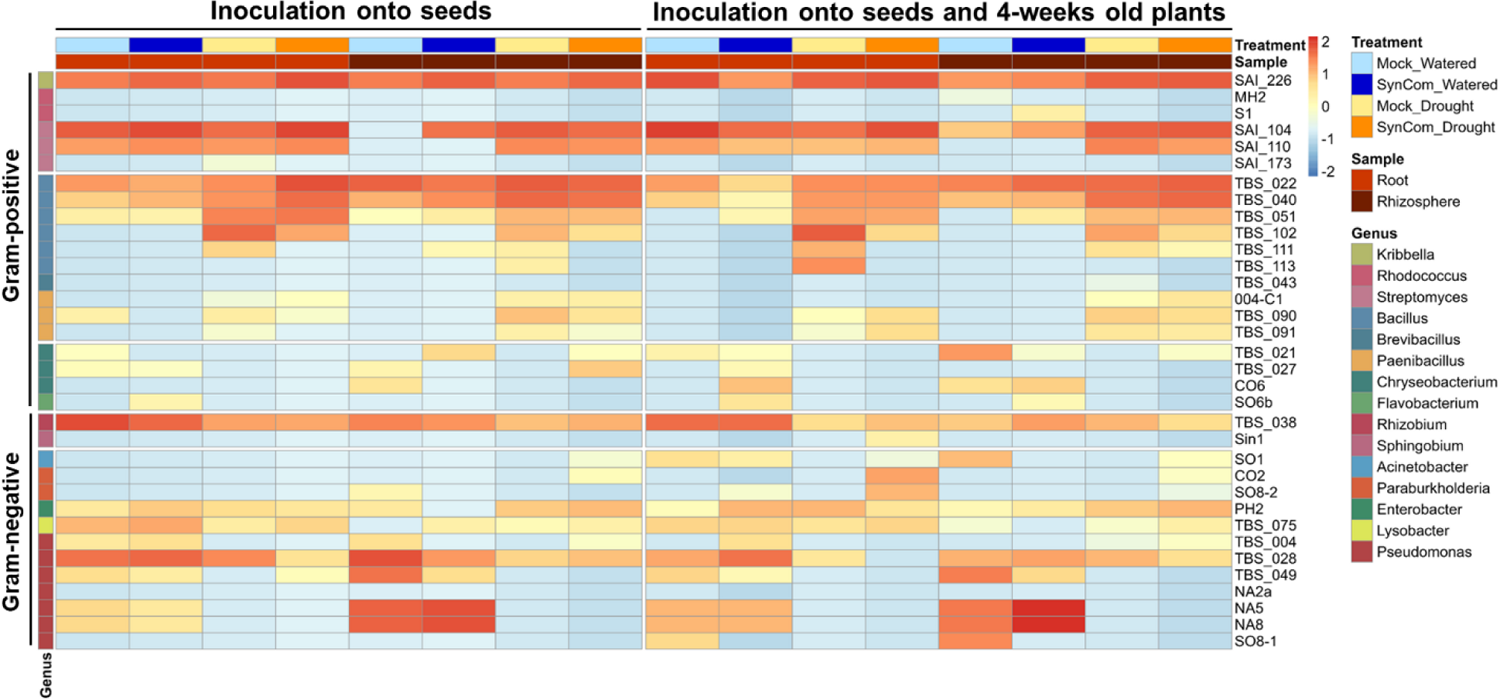
Long-term lack of engraftment of the dSynCom in the sorghum root microbiome under field conditions. Heat map presents the abundance of 16S rRNA ASVs identified to each dSynCom member across samples treated with mock or dSynCom once (seeds only) or twice (seeds and 4-weeks old plants). Sample types (roots in red and rhizosphere in dark red), inoculation (mock in light colors and dSynCom in dark colors), and water treatments (watered in blues and drought in browns) are shown at the top of the heat map. Genera and members included in the dSynCom are shown at the left and right sides, respectively. Abundance is presented in log2 scale of the sum of 20 samples per treatment (n=320 samples in total).

**Figure 7.**
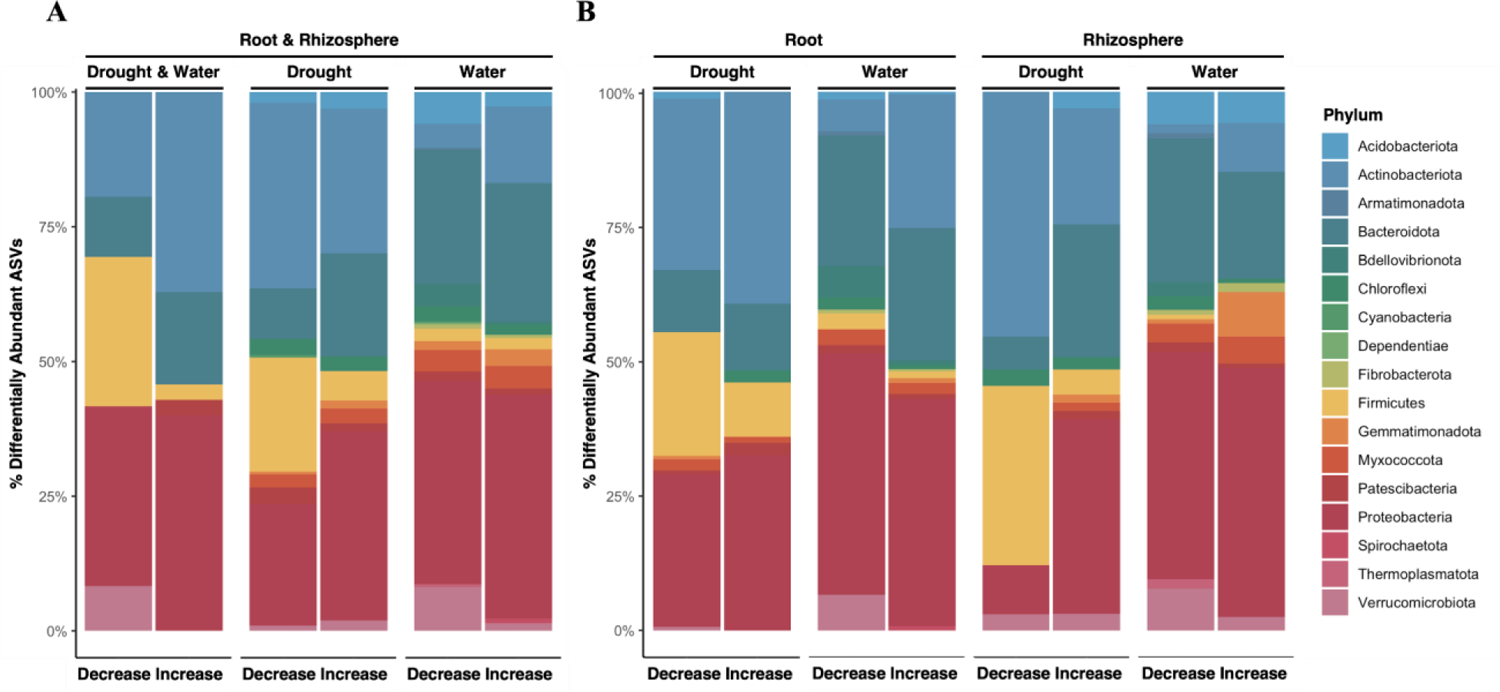
Application of dSynCom induces changes in community composition. A) Proportions of differentially abundant ASVs (log2FoldChange > 1.5, *P* < 0.05) either decreasing or increasing in response to dSynCom application including both root and rhizosphere compartments across both drought and watered conditions, drought only, and water only. B) Results when root and rhizosphere compartments are separately compared (n=320 samples in total).

**Figure 8.**
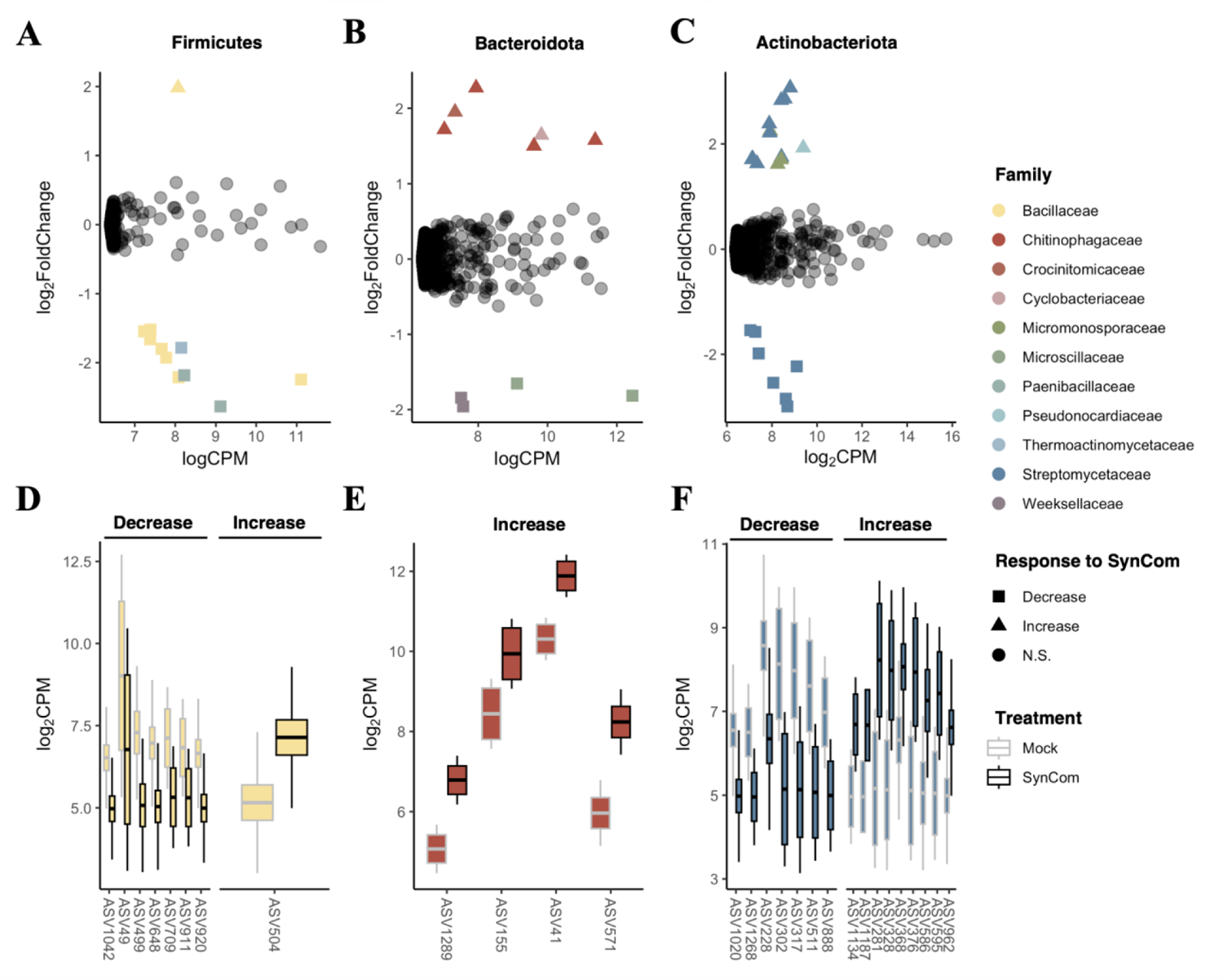
dSynCom impacts bacterial community member abundances in a lineage-dependent manner. Responses to dSynCom when comparing across all samples, including both root and rhizosphere samples, under both drought and water regime. LogCPM represents the mean abundance across both treatments. Negative and positive log2FoldChange values indicate decreased and increased abundance upon dSynCom treatment, respectively. Results separated by phylum (A-C). Significant (log2FoldChange > 1.5, *P* < 0.05) differentially abundant ASVs colored by family. Shape corresponds to changes in response to dSynCom application. Boxplots for notable family-wise trends (D-F) include divergent responses among *Streptomycetaceae*, increases in abundance in *Chitinophagaceae*, and decreases among *Bacillaceae*.

Despite a lack of evidence of dSynCom persistence within the root environment, positive impacts on plant phenotype were again observed following dSynCom treatment. As was observed previously in the lab-based *in planta* experiments, the shoot fresh weight, dry weight, and shoot length were all impacted. Specifically, each of these phenotypes was enhanced with the dSynCom treatment (both seed and 4-week old seedling inoculations) compared to mock-treated plants under both water and drought treatments (Figure 9A-D). In addition, we also observed a positive impact on the root phenotypes of watered plants that had been inoculated twice with the dSynCom (Supplemental Figure 10, Supplemental Tables 6-7), with significant effects in eight of the 36 features analyzed with the RhizoVision Crown system (Seethepalli et al., 2020). For root phenotypes, no significant effect was observed in either the shoot or root phenotypes on those plants that were inoculated only onto seeds (Supplemental Figure 10-11, Supplemental Tables 6-7). Interestingly, the number of tillers increased for both plants inoculated onto seeds or inoculated twice (seeds and seedlings) with the dSynCom under normal irrigation (Figure 9E, Supplemental Figure 11), which could partially explain overall increases in above ground yield and biomass. Finally, after a period of three months of restored irrigation (25 wap), the shoot fresh weight and dry weight were measured for all treatment groups; we observed increases in both phenotypes for plants inoculated onto seeds or twice with dSynCom (Figure 10A-B,D-E) under the watered treatment, although no significant effect was observed on droughted plants at this stage of development. Additionally, we noted that dSynCom treatment also positively impacted seed yield for plants under the normal irrigation treatment (Figure 10C,F). Altogether, these results demonstrate that treatment with the dSynCom is effective in increasing biomass and yield-related traits in the field as well as in the laboratory.

**Figure 9.**
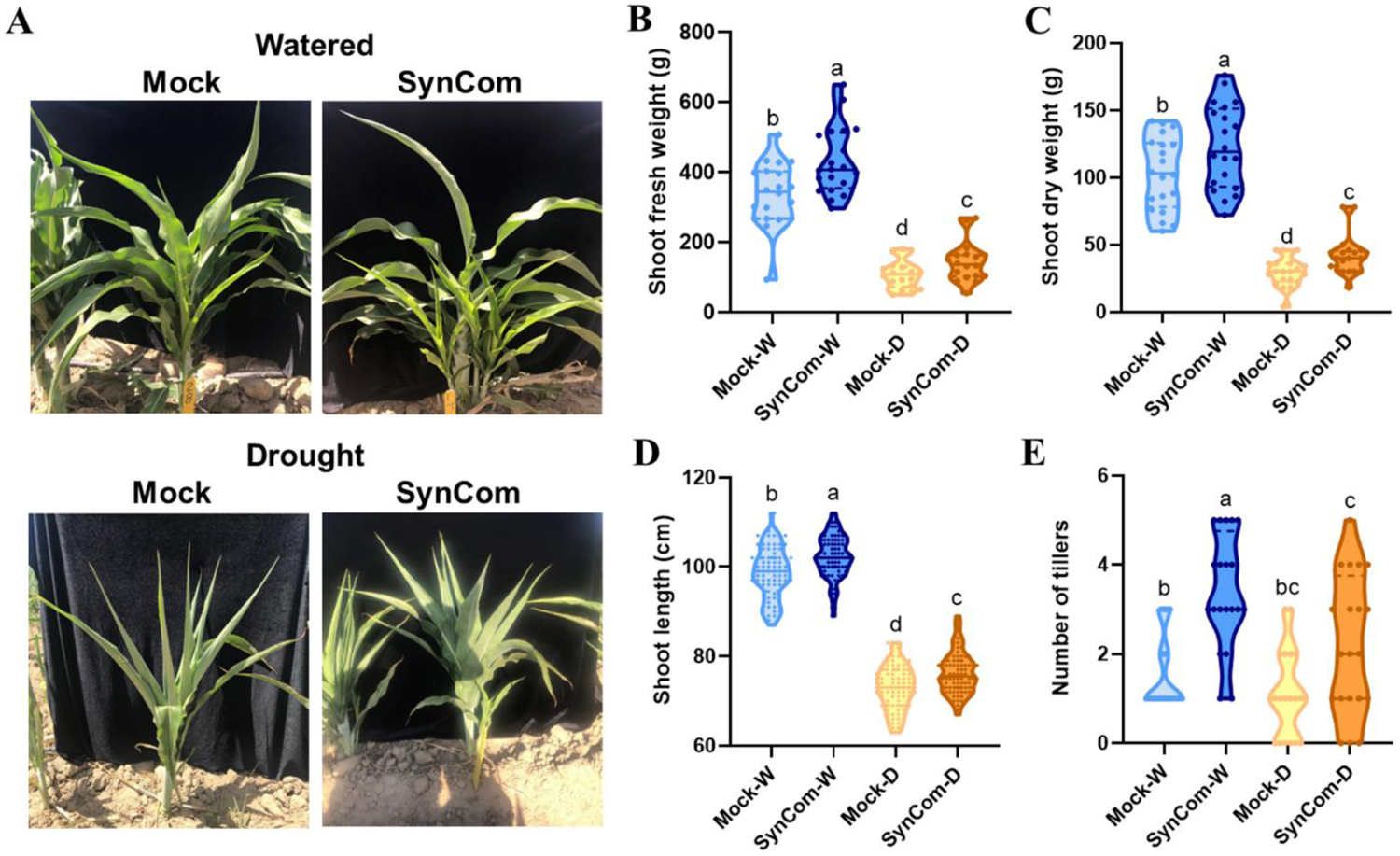
dSynCom enhances the shoot biomass of sorghum plants in the field at nine weeks after planting. A) Representative pictures of watered or drought stressed plants inoculated twice (seeds and 4-week old plants) with mock or dSynCom. B) Shoot fresh weight. C) Shoot dry weight. D) Shoot length. E) Tiller number. W=normal irrigation (blue colors); D=drought stress (brown colors). Letters above represent statistical differences among treatments by Brown-Forsythe and Welch ANOVA one-way, Holm-Šídák post hoc test *P* < 0.05 for A-B), ANOVA one-way, Tukey post hoc test *P* < 0.05 for D), and Kruskal-Wallis test, Dunn’s Test of multiple comparisons *P* < 0.05 for E). A total number of 80 plants were harvested, 20 plants per treatment.

**Figure 10.**
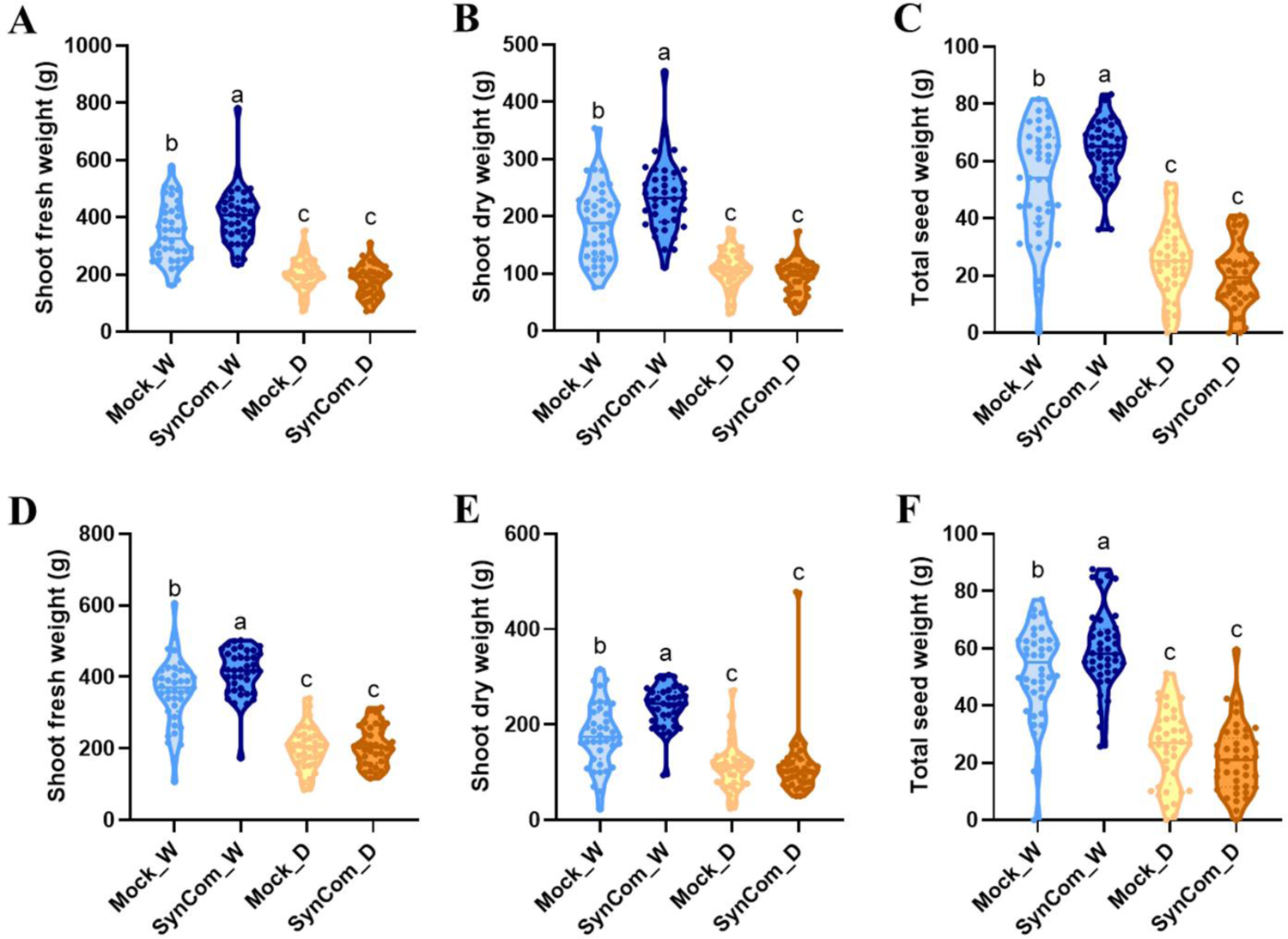
dSynCom improves seed production in the field. Shoot phenotyping of sorghum plants after three months of restored irrigation inoculated with the dSynCom once (A-C) and twice (D-F). A,D) Shoot fresh weight. B,E) Shoot dry weight. C,F) Total seed weight. W=normal irrigation (blue colors); D=drought stress (brown colors). Letters above represent statistical differences among treatments by Brown-Forsythe and Welch ANOVA one-way, Holm-Šídák post hoc test *P* < 0.05 for A-E), and ANOVA one-way, Tukey post hoc test *P* < 0.05 for F) of 317 plants harvested in total.

## DISCUSSION

In this study, a defined synthetic community was developed using a bottom-up approach with 16 genera representative of taxa commonly found within the rhizosphere microbiome of sorghum. One clear observation from our studies is that the community structure of the assembled dSynCom varied significantly depending on growth environment and the presence or absence of specific community members or additional native microbes. In plate-based assays, despite our efforts to include in the media complex metabolites typically provided by the plant, Gram-positive bacteria were less successful at persisting across multiple passages compared to their Gram-negative counterparts. This may in part be explained by differences in growth strategies of these two groups. Based on the r/K selection theory (Andrews and Harris, 1986), it has been proposed that Gram-negative bacteria commonly represent r-strategists while Gram-positive bacteria represent K-strategists (de Vries and Shade, 2013), where r-strategists favor resource-rich environments, and K-strategists are able to better adapted to crowded and resource-limited spaces (Yin et al., 2022). Both of the media used in this experiment, SSM and SSM-Ex, represent resource-rich environments full of compounds that can be rapidly metabolized. Similar results have been observed in other studies, where the *in vitro* influence of the nutritional content in the experimental environment has driven SynCom composition, with favor falling to Gram-negative bacteria when simple carbon sources are plentiful (Weiss et al., 2022). Alternatively, the failure of Gram-positives to thrive in our media-based growth environments could also be explained by moisture content; Gram-negative bacteria are typically adapted to low osmolarity environments with high water activity, while Gram-positive bacteria, by virtue of the strength of their cell walls, are capable of survival and growth in less dilute systems with lower water activity (Daniel et al., 2004). As compared to the majority of soils and rhizospheres, agar media provides a relatively high water activity environment, potentially favoring the growth of Gram-negative members of our dSynCom, although biases in DNA extraction may be present.

We also observed that the inclusion of sorghum-exudate utilizers as part of our SynCom formulation shaped dSynCom assembly on gnotobiotically-grown plants (no additional microbes present), with negative impacts on the abundance of both *Bacillus* and *Streptomyces*. One possible explanation for this could be direct antagonism. Two of the sorghum-exudate utilizing strains in our study belongs to the genus *Acinetobacter*, which has previously been shown to have the ability to kill Gram-positive bacteria via a T6SS-mediated antagonism (Le et al., 2021). Alternatively, this observation could be explained by strain-level competition for resources. Ongoing work from our group suggests that *Streptomyces* are good solubilizers of a wide variety of novel metabolites (Oda et al., 2023). We propose that *Streptomyces* may directly use sorghum-specific root exudate compounds such as sorgoleone and introduction of other utilizers creates competition for these carbon sources. Similar examples of substrate competition modulating microbiome composition have been observed in reduced-complexity microbial communities (Berry and Widder, 2014; Lozano et al., 2019; Weiss et al., 2022).

Not surprising given other results from this field of research, when the dSynCom was grown with the plant in the presence of native microbial diversity and the wider environmental heterogeneity typical of field environments, we were not able to detect long-term engraftment of any of the community members from our dSynCom formulation. Analogous results have been observed in many other systems where applied microbes that colonize well in gnotobiotic conditions or greenhouse settings did not effectively colonize the root microbiome in the field (Kaur et al., 2022; Liu et al., 2022; Schmitz et al., 2022). These types of results are often hypothesized to be the result of comparatively poorer fitness of the introduced strains for the specific environmental conditions present in the targeted environment. However, in our specific case all dSynCom members were originally cultured from sorghum roots or soils taken from this same field location, suggesting the likelihood of other factors at play. As an alternative explanation, root and rhizosphere microbiomes have been shown to be dynamic across plant development, with taxonomic variation occurring throughout the growing season (Gao et al., 2019). This could allow for the possibility that our dSynCom did show early engraftment with later replacement with alternative soil-derived taxa in response to plant metabolism, plant root development, or edaphic changes later in the experiment. In the future, broader longitudinal sampling of microbial community abundances would allow for reporting on early dSynCom colonization dynamics following initial inoculation.

Interestingly, despite a lack of evidence of long-term engraftment of the dSynCom in our field study, there was observable impact on the broader root microbiome composition. Other studies have also found that synthetic communities treatment can lead to rewiring of existing community structure without significant increases in the abundances of SynCom community members (Deng et al., 2019; Kaur et al., 2022; Liu et al., 2022). In our experiment, at the time of harvest (nine weeks after planting) we observed an enrichment in Bacteriodota phyla under drought conditions in plants that had received the inoculation compared to the mock control. In another study, it has been reported that increases in the abundances of this group of taxa was positively correlated with the total C, N and P in soil (Deng et al., 2020). In particular, members of the Bacteriodota family *Chitinophagacea*, which showed strong across the board increases following dSynCom treatment, can degrade complex organic matters such as chitin and cellulose (Rosenberg, 2014) and show β-glucosidase activity (Bailey et al., 2013) which can help to provide C and N to the plant. By contrast, the family *Bacillaceae* was mostly depleted in the dSynCom-inoculated plants under both irrigation treatments despite being one of the dominant taxonomic groups in our dSynCom formulation (n=18 *Bacillus* strains). One possible explanation for this observed microbiome restructuring might be the priming of immune responses and immune memory. Numerous studies have explored how plant defense signaling regulates bacterial colonization in plants (Liu et al., 2017) with the activation of salicylic acid (SA) and jasmonic acid (JA)/ethylene (ET) signaling pathways suppressing the colonization of specific bacterial lineages in a variety of plant systems (Iniguez et al., 2005; Miché et al., 2006; Kniskern et al., 2007). For example, Jiang et al., (2016) demonstrated that application of a *Bacillus cereus* strain elicits the Induced Systemic Resistance (ISR) by simultaneously activating both the SA and JA/ET signaling pathways. This hormonal activation leads to modulation of the endophytic bacterial community in Arabidopsis, with primary impacts observed on other *Bacillus* species (Kniskern et al., 2007). Similarly, another study of plant ISR has shown that priming with a pathogen early in development can lead to increased resistance to colonization by all members belonging to that bacterial group later in development (Nguyen et al., 2020).

Equally interesting to this modulation of the native root microbiome is the clear impact of dSynCom treatment on plant phenotype despite a lack of clear engraftment at harvest. These improvements impacted both above and below ground fitness, and for some phenotypes of dSynCom treated plants, overall shoot biomass, seed production, and root biomass all showed significant increases independently of the watering treatment. While typically there are tradeoffs for growth promotion where benefit is observed in a specific tissue type (García et al., 2020), there are also cases in which all plant parts show increases following microbial inoculation (Dey et al., 2004; Egamberdiyeva, 2007; Ferreira et al., 2013). Our mechanistic understanding of what leads to these universally positive responses remains limited, but one possibility involves phytohormones. Regulation of the shoot/root growth ratio is a conserved developmental mechanism in plants, which guarantees the highest progeny production under environmental changes (Kurepa and Smalle, 2022). The hormones auxin and cytokinin regulate the shoot/root growth ratio in which cytokinin promotes shoot and inhibits root growth, whereas auxin does the opposite (Skoog and Miller, 1957; Li et al., 2009; Kurepa et al., 2014). Mounting evidence has demonstrated that numerous microorganisms have been shown to have the ability to manipulate levels of these hormones in and around plant tissues (Hacquard et al., 2015) leading to more robust phenotypes. Alternatively, it has recently been suggested that epigenetics may play a role in the long-term modulation of plant phenotype in response to microbial amendments. In a long-term experiment involving pokeweed, plant growth promotion was observed alongside modifications to DNA methylation in the plant genome long after elimination of the inoculum from the native microbiome (Chen et al., 2022). Future work to investigate the impact of our dSynCom treatment on plant hormones, nutrient use pathways, and epigenetic states may help clarify the mode of action by which our dSynCom increases plant fitness.

Finally, the results from our study and other recently published work (Deng et al., 2019) suggest it may be helpful to distinguish between cases in which persistence of a microbial inoculum is likely to be required *versus* helpful. In the context of our field trial, persistence of the members of the dSynCom did not appear to be strictly required to impact the plant performance. However, in other scenarios, persistence may be a prerequisite for success. For instance, in the case of microbially-mediated soil remediation, microbial presence may be necessary for long-term services it provides. Phytoremediation studies of diesel-contaminated soils have shown a positive correlation between the persistence of the population of the inoculated microbes in the rhizosphere and endosphere of the host plant and the improvement of plant growth, hydrocarbon degradation, and toxicity reduction (Afzal et al., 2013). Similarly, for the long-term prevention of a specific pathogen agent through biocontrol, short term engraftment may not be sufficient for prevention. The mode of action of many of these disease suppressive agents is often direct microbe-microbe interaction, principally mediated by allelochemicals including siderophores, antibiotics, cell wall-degrading enzymes, volatile organic compounds, alkaloids, and many other compounds (Oukala et al., 2021). As an example, Sheoran et al., (2015) demonstrated that black pepper-associated endophytic *Pseudomonas putida* BP25 could inhibit, by volatile emission, the proliferation of different pathogens including fungi and oomycetes, and plant-parasitic nematode. We propose that while a better understanding of the forces that act to limit the persistence of introduced microbes is critical to designing ones that can persist, it is also worth considering whether persistence is in fact required for the desired outcome.

## CONCLUSION

Interest in the use of microbial amendments in agriculture and the environment to achieve specific goals is increasing. Consistent challenges in achieving desired outcomes following deployment are often attributed to failure of these products to compete and engraft in their environment. In this study we simultaneously demonstrate that: 1) engraftment operates as a function of the microbial diversity in the system and growth environment parameters; and 2) that engraftment may not be strictly necessary over long periods of time for realization of specific outcomes. Collectively, these results point to the promise of syncom-based strategies for field deployment and use in improving agricultural activity.

## Data availability

All datasets and scripts for analysis are available through GitHub (https://github.com/CitlaliFG/dSynCom-PerCon) and all short read data can be accessed through NCBI BioProject xxx.

## Acknowledgements

We thank for their help in sample collection and field preparation the staff of the Kearney Agricultural Research Center. We also thank Claudia Castro for her constructive revision of this manuscript.

