## Supplemental material for "Defined synthetic microbial communities colonize and benefit field-grown sorghum"

### Supplemental figures

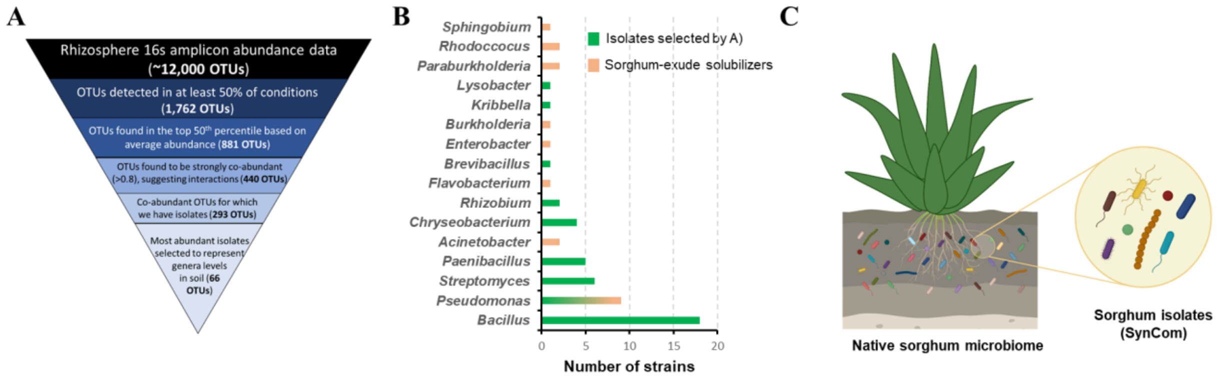

**Supplemental Figure 1.** dSynCom design using a bottom-up approach. A) Amplicon and network analysis-based selection of species for dSynCom development. B) Genera included in the dSynCom of the members selected based on the network analysis and/or sorghum-exudate solubilizers. C) Schematic representation of the member strains isolated from rhizosphere and roots of field grown Sorghum.

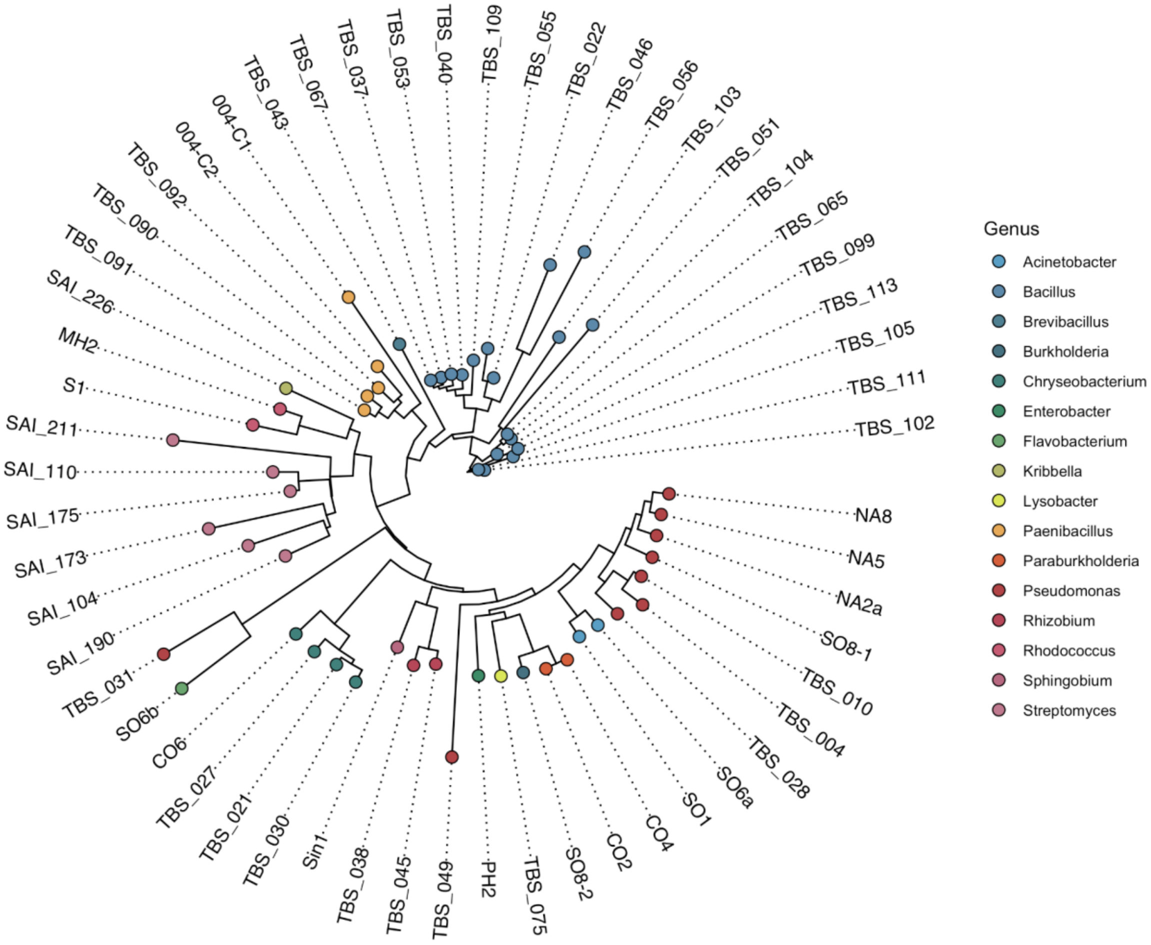

**Supplemental Figure 2.** Neighbor-joining tree of DSynCom members built from ClustalOmega multiple sequence alignment using Sanger sequenced 16S rRNA DNA sequences (~1000 bp length) and colored by genus.

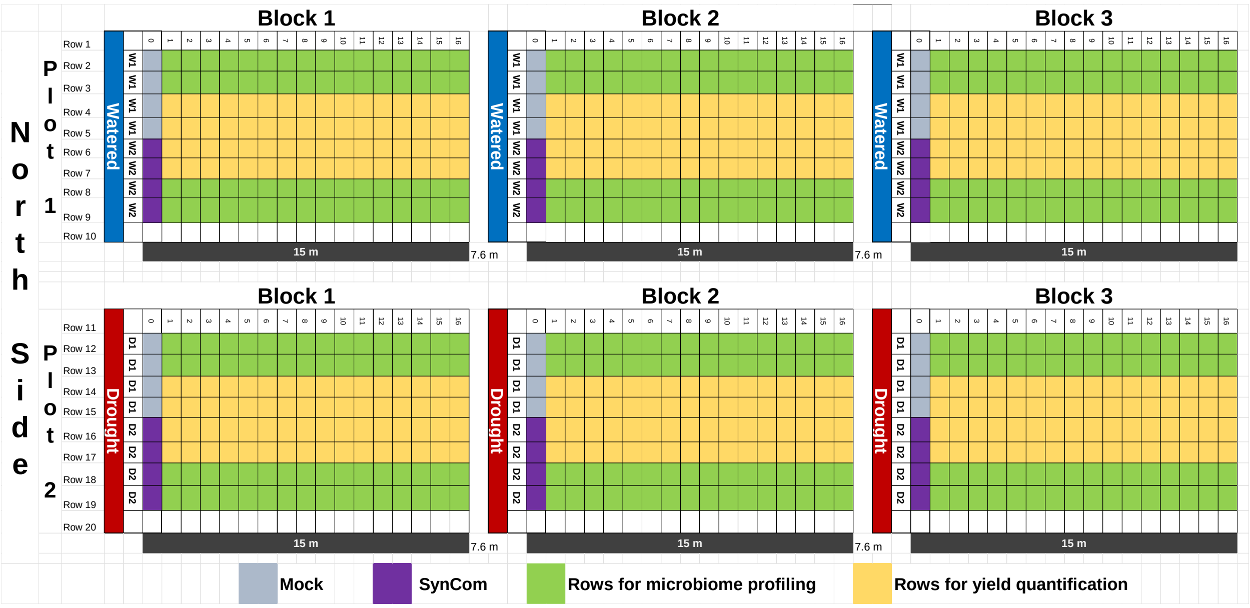

**Supplemental Figure 3.** Schematic diagram of the field layout. Six blocks representing two water treatments (blue: watered and red: drought), two inoculation treatments (gray: mock and purple: dSynCom), and three replicates were arranged in three blocks across a 60 m by 30 m field, with 7.6 m spacing between plots. Each plot consisted of ten 15m long rows, each containing approximately 200 plants spaced 10cm apart. Plants inoculated twice (seeds and 4-weeks old plants) with mock or dSynCom were labbed with a numerated stake. Samples for microbiome profiling and plant phynotyping at 9 wap were collected from green rows and yellow rows were reserved for yield measurements at 25 wap.

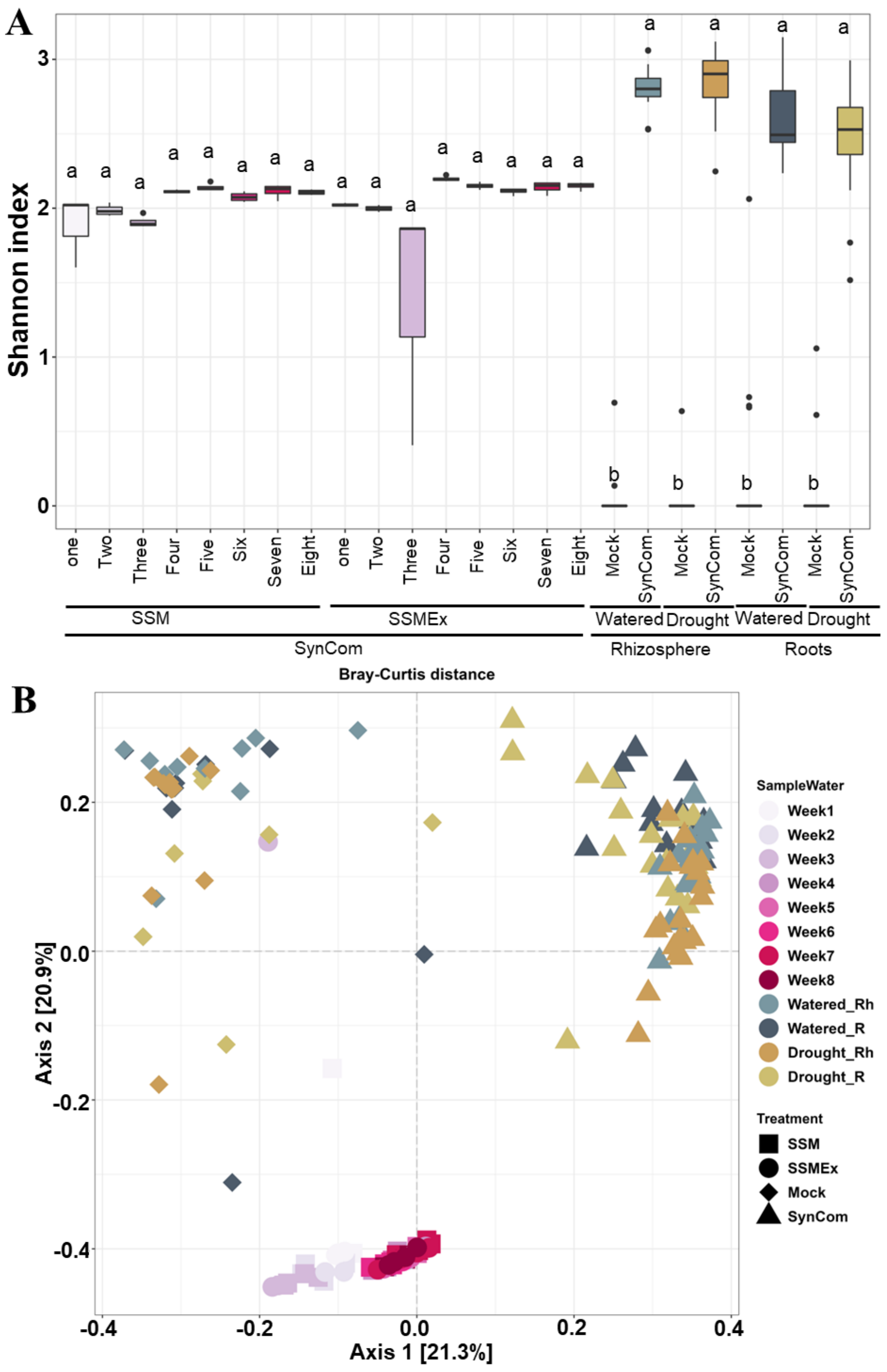

**Supplemental Figure 4**. 16S rRNA profiling of the dSynCom on agar plates and lab-based *in planta* experiments. A) Alpha diversity using the Shannon index. B) Beta diversity analysis using Bray-Curtis distance in a Principal Components Analysis. SSM= Synthetic Soil Media; SSM-Ex SSM + sorghum exudates; 1-8= weeks of passages on agar plates; Rh=Rhizosphere; R= Roots. Letters above in A) represent statistical differences among treatments by Kruskal-Wallis test, Dunn's Test of multiple comparisons p<0.05 of 328 samples in total.

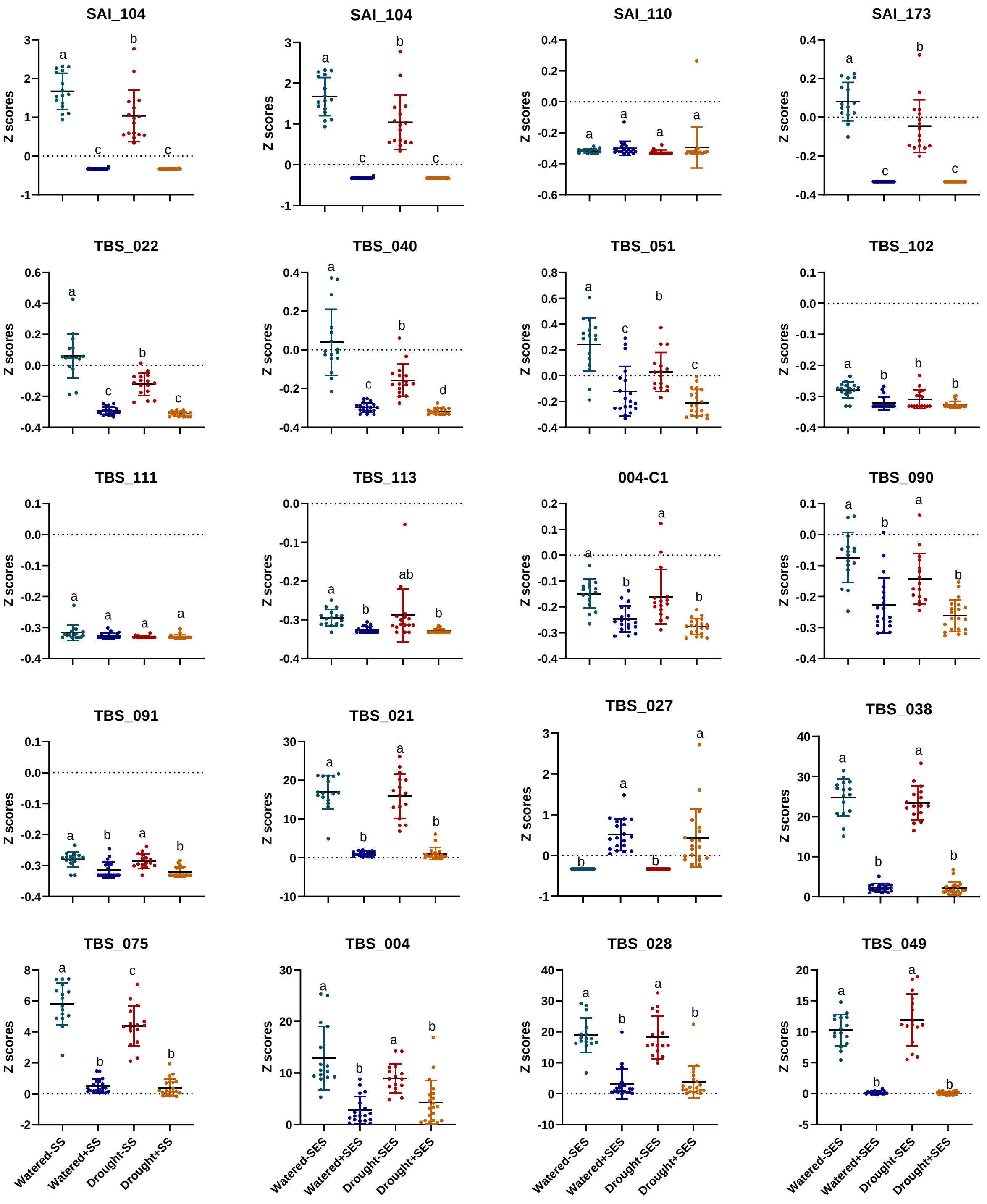

**Supplemental Figure 5.** Abundance of the dSynCom members identified in lab-based *in planta* experiments. The abundance is presented in Z cores across treatments, watered with (blue) and without (green) sorghum-exudate solubilizers (SES), drought with (red) and without (orange) sorghum-exudate solubilizers. Letters above represent statistical differences among treatments by Brown-Forsythe and Welch ANOVA one-way, Holm-Šídák post hoc test p<0.05 of 218 samples in total.

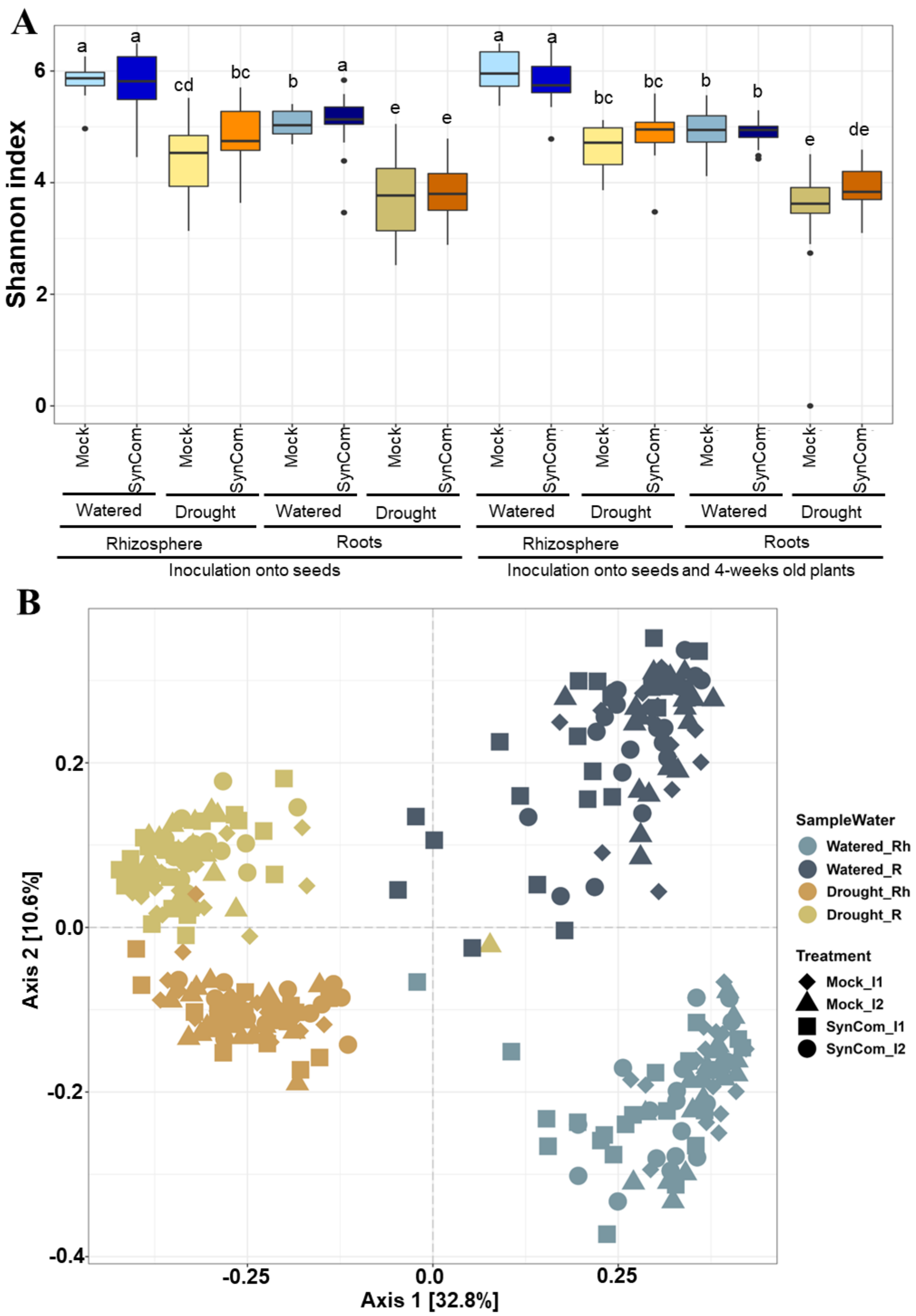

**Supplemental Figure 6.** 16S rRNA profiling of the sorghum microbiome of the field trial. A) Alpha diversity using the Shannon index. B) Beta diversity analysis using Bray-Curtis distance in a Principal Components Analysis. Letters above in A) represent statistical differences among treatments by ANOVA one-way test and Tukey post hoc test with $p \leq 0.05$ of 320 samples in total. Abbreviations in B): Rh= Rhizosphere; R= Roots; I1= inoculated once (seeds); I2= inoculated twice (seeds and 4-weeks old plants).

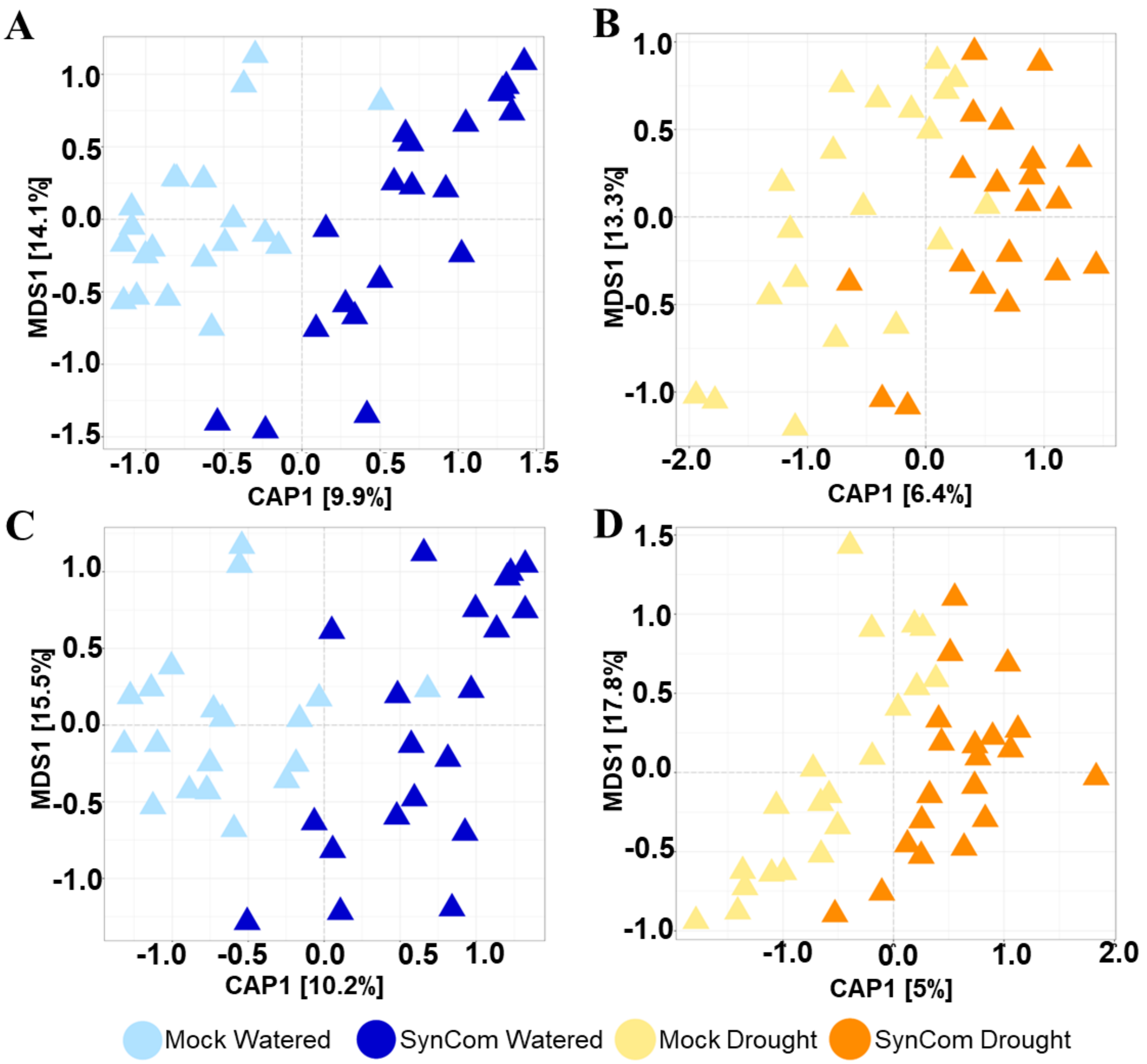

**Supplemental Figure 7.** Beta diversity analysis of the Sorghum microbiome of the field trial. Constrained Analysis of Principal Coordinates (CAP) ordination plots of the microbial community in the sorghum (A-B) rhizosphere and (C-D) roots samples under (A,C) normal irrigation (blue colors) and (B,D) drought stress (brown colors) inoculated onto seeds with Mock or dSynCom.

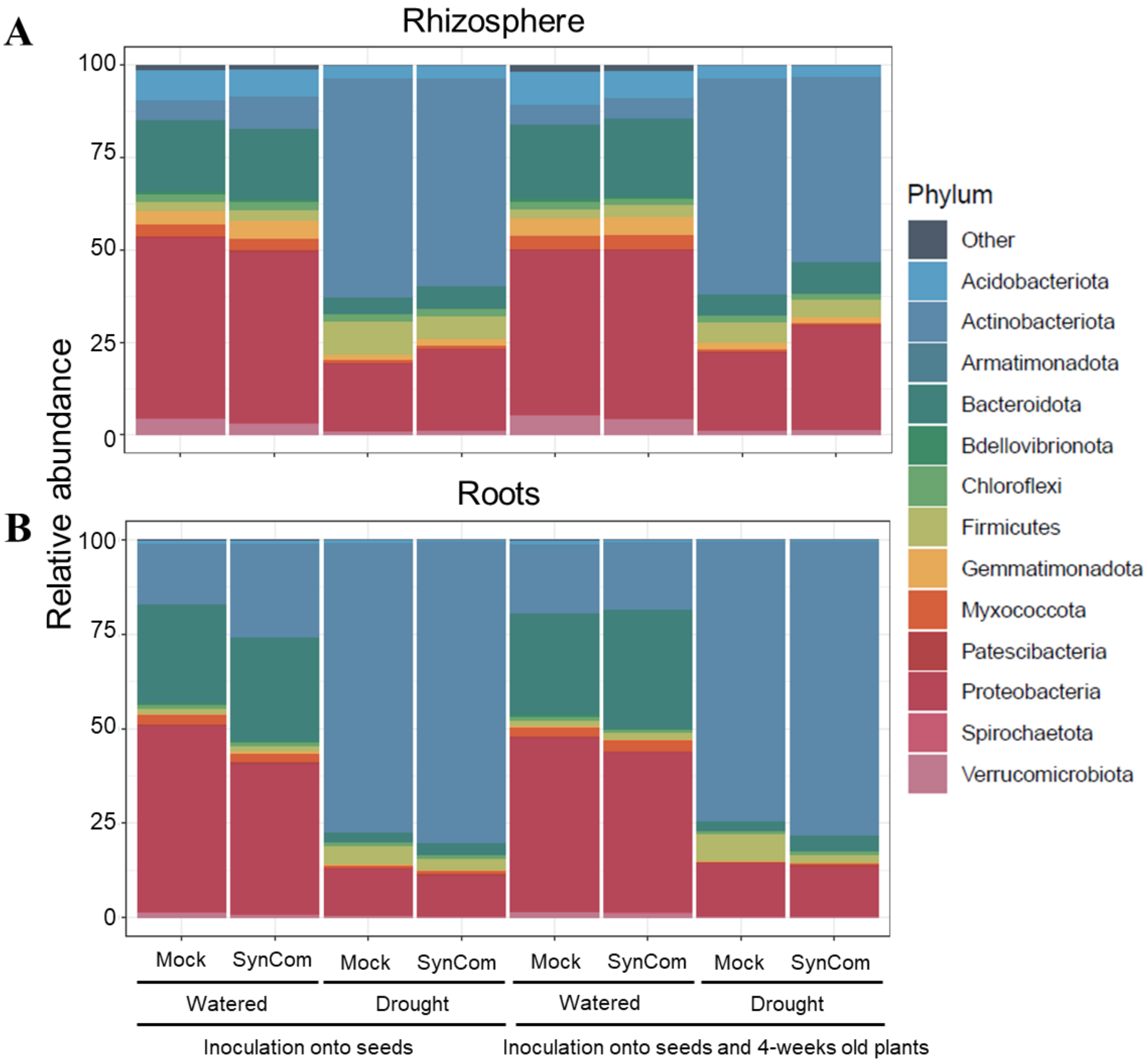

**Supplemental Figure 8.** Relative abundance plots for the most abundant bacterial phyla of the field trial. Percent relative abundance of the top 13 most abundant phyla for watered and droughted stress plants inoculated once (seeds) or twice (seeds and 4-weeks old plants) in rhizosphere (A) and roots (B).

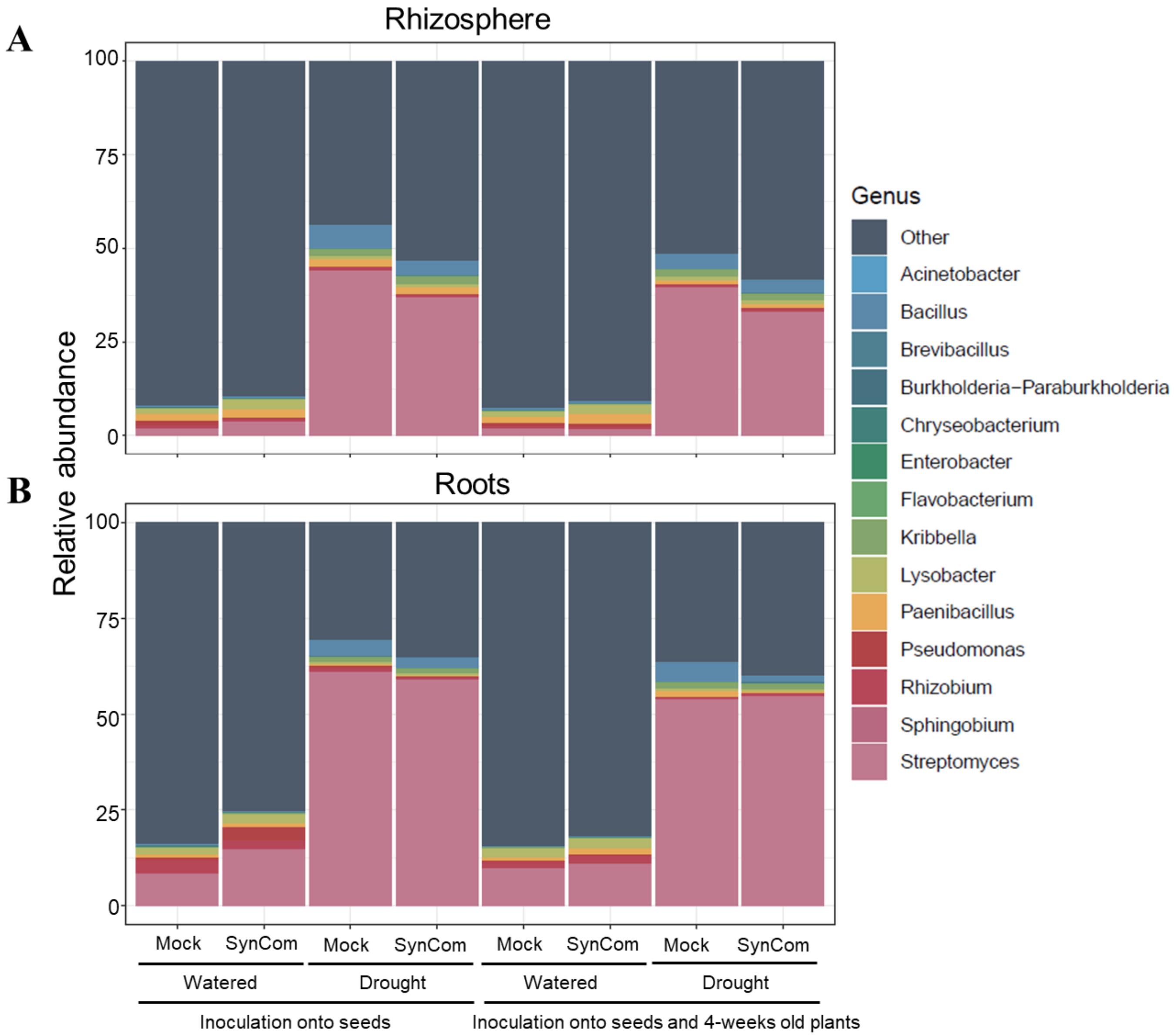

**Supplemental Figure 9.** Relative abundance plots for the bacterial genera of the field trial. Percent relative abundance of the genera present in the dSynCom for watered and droughted stress plants inoculated once (seeds) or twice (seeds and 4-weeks old plants) in rhizosphere (A) and roots (B) of the field trial.

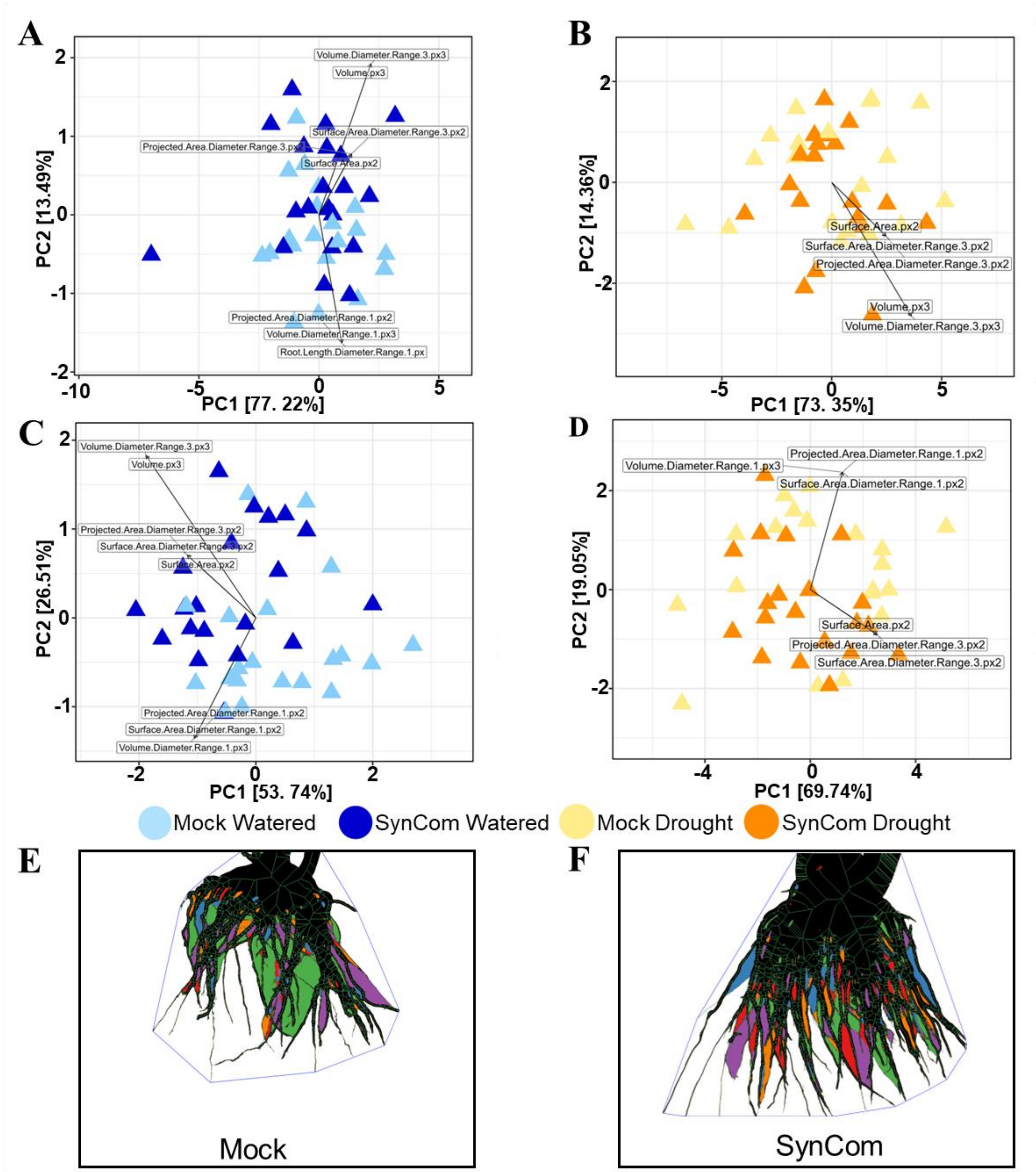

**Supplemental Figure 10.** Root phenotyping of sorghum plants of the field trial at ten weeks after planting. Principal Component Analysis (PCA) plots of roots samples under (A,C) normal irrigation (blue colors) and (B,D) drought stress (brown colors) inoculated onto seeds (A-B) or onto seeds and 4-weeks old plants (C-D) with mock or dSynCom. Representative images of roots treated with mock (E) or dSynCom (F) digitized using RhizoVision Crown system (Seethepalli et al., 2020).

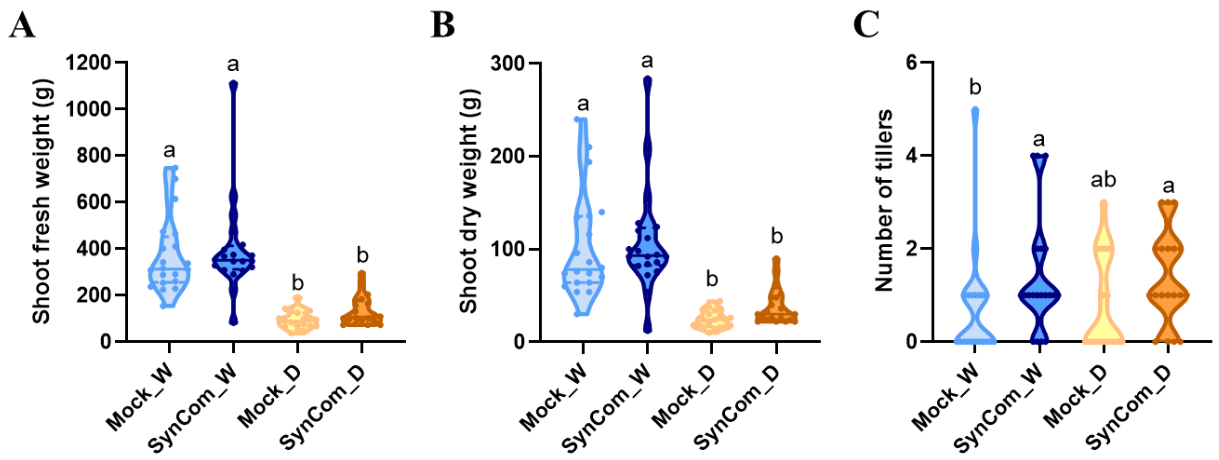

**Supplemental Figure 11.** Shoot phenotyping of sorghum plants of the field trial at 9 wap with seeds inoculated with mock or dSynCom. A) Shoot fresh weight, B) Shoot dry weight. C) Tillers number. W= normal irrigation (blue colors); D= Drought stress (brown colors). Letters above represent statistical differences among treatments by Kruskal-Wallis test, Dunn's Test of multiple comparisons $p<0.05$ of 80 harvested plants, 20 plants per treatment.

**Supplemental Tables**

**Supplemental Table 1.** Strains members of the dSynCom used in this study.

| dSynCom Strain # | Strain Label | Genus | Species | Evidence for taxonomic assignment | Skills | Plating schedule [1] | ASV group detected in 16S rRNA data | ID indicated in heat maps |
|---|---|---|---|---|---|---|---|---|
| 1 | SAI_226 | *Kribbella* | *shirazensis* | | network-based analysis | Day 1 | SAI_226 | SAI_226 |
| 2 | 004-C2 | *Paenibacillus* | *favisporus* | | network-based analysis | Day 2 | 004-C1/004-C2 | 004-C1 |
| 3 | SAI_104 | *Streptomyces* | *ambofaciens* | | network-based analysis | Day 2 | SAI_104 | SAI_104 |
| 4 | SAI_110 | *Streptomyces* | *graminofaciens* | | network-based analysis | Day 2 | SAI_110/SAI_175/SAI_211 | SAI_110 |
| 5 | SAI_173 | *Streptomyces* | *albogriseolus* | | network-based analysis | Day 2 | SAI_173/SAI_190 | SAI_173 |
| 6 | SAI_175 | *Streptomyces* | *graminofaciens* | | network-based analysis | Day 2 | SAI_110/SAI_175/SAI_211 | SAI_110 |
| 7 | SAI_190 | *Streptomyces* | *albogriseolus* | | network-based analysis | Day 2 | SAI_173/SAI_190 | SAI_173 |
| 8 | SAI_211 | *Streptomyces* | *graminofaciens* | | network-based analysis | Day 2 | SAI_110/SAI_175/SAI_211 | SAI_110 |
| 9 | MH2 | *Rhodococcus* | *aetherivorans* | *GTDBtk* | MHPP solubilizer | Day 2 | MH2 | MH2 |
| 10 | SO8-2 | *Burkholderia* | *anthina_A* | *GTDBtk* | sorgoleone | Day 2 | SO8-2 | SO8-2 |

| | | | | | | | | |
|---|---|---|---|---|---|---|---|---|
| | | | | | solubilizer | | | |
| 11 | CO6 | *Chryseobacterium* | *geocarphospharae* | *GTDBtk* | o-coumaric acid solubilizer | Day 2 | CO6 | CO6 |
| 12 | PH2 | *Enterobacter* | *ludwigii* | | o-coumaric acid solubilizer | Day 2 | PH2 | PH2 |
| 13 | CO2 | *Paraburkholderia* | *sp.* | *GTDBtk* | o-coumaric acid solubilizer | Day 2 | CO2/CO4 | CO2 |
| 14 | S1 | *Rhodoccocus* | *rhodochrous* | | Pipecolic acid solubilizer | Day 2 | S1 | S1 |
| 15 | TBS_038 | *Rhizobium* | *pusense* | | network-based analysis | Day 3 | TBS_038/TBS_045 | TBS_038 |
| 16 | TBS_045 | *Rhizobium* | *pusense* | | network-based analysis | Day 3 | TBS_038/TBS_045 | TBS_038 |
| 17 | TBS_075 | *Lysobacter* | *soli* | | network-based analysis | Day 3 | TBS_075 | TBS_075 |
| 18 | 004-C1 | *Paenibacillus* | *lautus* | | network-based analysis | Day 4 | 004-C1/004-C2 | 004-C1 |
| 19 | TBS_090 | *Paenibacillus* | *lautus* | | network-based analysis | Day 4 | TBS_090/TBS_092 | TBS_090 |
| 20 | TBS_091 | *Paenibacillus* | *lautus* | | network-based analysis | Day 4 | TBS_091 | TBS_091 |

| | | | | | | | | |
|---|---|---|---|---|---|---|---|---|
| 21 | TBS_092 | *Paenibacillus* | *lautus* | | network-based analysis | Day 4 | TBS_090/TBS_092 | TBS_090 |
| 22 | Sin1 | *Sphingobium* | *sp.* | *GTDBtk* | sorgoleone solubilizer | Day 4 | Sin1 | Sin1 |
| 23 | CO4 | *Paraburkholderia* | *sp.* | *GTDBtk* | o-coumaric acid solubilizer | Day 4 | CO2/CO4 | CO2 |
| 24 | NA2a | *Pseudomonas* | | *GTDBtk* | sorgoleone solubilizer | Day 4 | NA2 | NA2 |
| 25 | SO1 | *Acinetobacter* | *pittii* | *GTDBtk* | sorgoleone solubilizer | Day 4 | S01/SO6a | S01 |
| 26 | SO6a | *Acinetobacter* | *sp.* | *GTDBtk* | sorgoleone solubilizer | Day 4 | S01/SO6a | S01 |
| 27 | SO6b | *Flavobacterium* | *sp.* | *GTDBtk* | sorgoleone solubilizer | Day 4 | SO6b | SO6b |
| 28 | NA5 | *Pseudomonas* | *entomophila* | | sorgoleone solubilizer | Day 4 | NA5 | NA5 |
| 29 | SO8-1 | *Pseudomonas_E* | *sp.* | *GTDBtk* | sorgoleone solubilizer | Day 4 | SO8-1 | SO8-1 |
| 30 | NA8 | *Pseudomonas* | *entomophila* | | sorgoleone solubilizer | Day 4 | NA8 | NA8 |

| | | | | | | | | |
|---|---|---|---|---|---|---|---|---|
| 31 | TBS_004 | *Pseudomonas* | *frederiksbergensis_A* | *GTDBtk* | network-based analysis | Day 4 | TBS_004/TBS_010 | TBS_004 |
| 32 | TBS_010 | *Pseudomonas* | *frederiksbergensis_A* | *GTDBtk* | network-based analysis | Day 4 | TBS_004/TBS_010 | TBS_004 |
| 33 | TBS_021 | *Chryseobacterium* | *gleum* | | network-based analysis | Day 4 | TBS_021/TBS_030 | TBS_021 |
| 34 | TBS_022 | *Bacillus* | *zanthoxyli* | | network-based analysis | Day 4 | TBS_022/TBS_037/TBS_053/TBS_056 | TBS_022 |
| 35 | TBS_027 | *Chryseobacterium* | *proteolyticum* | | network-based analysis | Day 4 | TBS_027 | TBS_027 |
| 36 | TBS_028 | *Pseudomonas_E* | *sp900156465* | *GTDBtk* | network-based analysis | Day 4 | TBS_028/TBS_031 | TBS_028 |
| 37 | TBS_030 | *Chryseobacterium* | *gleum* | | network-based analysis | Day 4 | TBS_021/TBS_030 | TBS_021 |
| 38 | TBS_031 | *Pseudomonas_E* | *sp900156465* | *GTDBtk* | network-based analysis | Day 4 | TBS_028/TBS_031 | TBS_028 |
| 39 | TBS_037 | *Bacillus* | *megaterium* | | network-based analysis | Day 4 | TBS_022/TBS_037/TBS_053/TBS_056 | TBS_022 |
| 40 | TBS_040 | *Bacillus* | *zanthoxyli* | | network-based analysis | Day 4 | TBS_040/TBS_046/TBS_055/TBS_067/1TBS_109 | TBS_040 |
| 41 | TBS_043 | *Brevibacillus* | *agri* | | network-based analysis | Day 4 | No found it | No found it |

| | | | | | | | | |
|---|---|---|---|---|---|---|---|---|
| 42 | TBS_046 | *Bacillus* | *zanthoxyli* | | network-based analysis | Day 4 | TBS_040/TBS_046/TBS_055/TBS_067/1TBS_109 | TBS_040 |
| 43 | TBS_049 | *Pseudomonas_E* | *pudica* | *GTDBtk* | network-based analysis | Day 4 | TBS_049 | TBS_049 |
| 44 | TBS_051 | *Bacillus* | *pumilus* | | network-based analysis | Day 4 | TBS_051/TBS_065/TBS_099/TBS_103/TBS_104 | TBS_051 |
| 45 | TBS_053 | *Bacillus* | *megaterium* | | network-based analysis | Day 4 | TBS_022/TBS_037/TBS_053/TBS_056 | TBS_022 |
| 46 | TBS_055 | *Bacillus* | *zanthoxyli* | | network-based analysis | Day 4 | TBS_040/TBS_046/TBS_055/TBS_067/1TBS_109 | TBS_040 |
| 47 | TBS_056 | *Bacillus* | *zanthoxyli* | | network-based analysis | Day 4 | TBS_022/TBS_037/TBS_053/TBS_056 | TBS_022 |
| 48 | TBS_065 | *Bacillus* | *pumilus* | | network-based analysis | Day 4 | TBS_051/TBS_065/TBS_099/TBS_103/TBS_104 | TBS_051 |
| 49 | TBS_067 | *Bacillus* | *aryabhattai* | | network-based analysis | Day 4 | TBS_040/TBS_046/TBS_055/TBS_067/1TBS_109 | TBS_040 |
| 50 | TBS_099 | *Bacillus* | *pumilus* | | network-based analysis | Day 4 | TBS_051/TBS_065/TBS_099/TBS_103/TBS_104 | TBS_051 |
| 51 | TBS_102 | *Bacillus* | *subtilis* | | network-based analysis | Day 4 | TBS_102/TBS_105 | TBS_102 |

| | | | | | | | | |
|---|---|---|---|---|---|---|---|---|
| 52 | TBS_103 | *Bacillus* | *pumilus* | | network-based analysis | Day 4 | TBS_051/TBS_065/TBS_099/TBS_103/TBS_104 | TBS_051 |
| 53 | TBS_104 | *Bacillus* | *pumilus* | | network-based analysis | Day 4 | TBS_051/TBS_065/TBS_099/TBS_103/TBS_104 | TBS_051 |
| 54 | TBS_105 | *Bacillus* | *subtilis* | | network-based analysis | Day 4 | TBS_102/TBS_105 | TBS_102 |
| 55 | TBS_111 | *Bacillus* | *subtilis* | | network-based analysis | Day 4 | TBS_111 | TBS_111 |
| 56 | TBS_113 | *Bacillus* | *haynesii* | | network-based analysis | Day 4 | TBS_113 | TBS_113 |
| 57 | TBS_109 | *Bacillus* | *zanthoxyli* | | network-based analysis | Day 4 | TBS_040/TBS_046/TBS_055/TBS_067/1TBS_109 | TBS_040 |

[1] Days before dSynCom preparation.

**Supplemental Table 2.** Recipe for MME and SSM. SSM is made using the MME media as a base and adding the nutrient sources listed here.

| MME Ingredients | | |
|---|---|---|
| **Ingredient** | **Final Amount** | **Unit** |
| $K_2HPO_4$ | 9.2 | mM |
| MOPS media | 20 | mM |
| NaCl | 4.3 | mM |
| $MgSO_4$ – 7 $H_20$ | 0.41 | mM |
| $CaCl_2$ – 2 $H_2O$ | 0.068 | mM |
| MME Trace Minerals | 1 | ml/L |
| **SSM Ingredients** | | |
| **Ingredient** | **Final Amount** | **Unit** |
| betaine hydrochloride | 2.147 | mM |
| sucrose | 2 | mM |
| L-asparagine monohydrate | 1.107 | mM |
| choline hydrochloride | 1.069 | mM |
| L-proline | 1.017 | mM |
| 4-hydroxybenzoate | 0.537 | mM |
| adenosine | 0.442 | mM |
| sn-glycero-3-phosphocholine | 0.409 | mM |
| L-valine | 0.29 | mM |
| L-leucine | 0.279 | mM |
| L-glutamine | 0.19 | mM |
| L-isoleucine | 0.188 | mM |
| allantoin | 0.163 | mM |
| D-pipecolic acid | 0.153 | mM |
| L-carnitine hydrochloride | 0.151 | mM |
| uridine | 0.121 | mM |
| D-galactitol / D-dulcitol | 0.102 | mM |
| D-mannitol | 0.102 | mM |
| L-arginine monohydrochloride | 0.099 | mM |
| p-coumaric acid (4-hydroxycinnamic acid) | 0.095 | mM |
| **1000X Trace Mineral Solution** | | |
| **Ingredient** | **Final Amount** | **Unit** |
| HCl | 1 | mL/L |
| $Na_4EDTA \times xH_2O$ | 380.17 | mM |
| $FeCl_3$ | 12.2 | mM |
| $H_3BO_3$ | 0.808 | mM |

| | | |
|---|---|---|
| $ZnCl_2$ | 0.367 | mM |
| $CuCl_2 \times 2H_2O$ | 0.176 | mM |
| $MnCl_2 \times 4H_2O$ | 0.253 | mM |
| $(NH_4)2MoO_4$ | 0.255 | mM |
| $CoCl_2 \times 6H_2O$ | 0.210 | mM |
| $NiCl_2 \times 6H_2O$ | 0.210 | mM |

**Supplemental Table 3.** PERMANOVA analysis of *in vitro* and *in planta* experiments 16S rRNA data with sorghum-exudate solubilizers included. PERMANOVA analysis was conducted using the Bray-Curtis distance of mock and dSynCom samples per media and water treatment.

| | **Factor** | **DF[1]** | **SumsOfSqs[2]** | **MeanSqs[3]** | **F Model** | **R2** | **Adjusted p-value[4]** |
|---|---|---|---|---|---|---|---|
| **Global** | Treatment | 3 | 28.041 | 9.347 | 43.590 | 0.387 | 0.001 |
| | SampletypeWeek | 8 | 1.836 | 0.230 | 1.070 | 0.025 | 0.343 |
| | WaterMedia | 1 | 0.352 | 0.352 | 1.641 | 0.005 | 0.103 |
| | Group | 11 | 2.008 | 0.183 | 0.851 | 0.028 | 0.878 |
| | Residuals | 188 | 40.312 | 0.214 | | 0.556 | |
| | Total | 211 | 72.550 | | | 1 | |
| ***In vitro*** | Media | 1 | 0.053 | 0.053 | 2.163 | 0.024 | 0.027 |
| | Week | 7 | 0.886 | 0.127 | 5.185 | 0.399 | 0.001 |
| | Group | 7 | 0.257 | 0.037 | 1.506 | 0.116 | 0.059 |
| | Residuals | 42 | 1.025 | 0.024 | | 0.462 | |
| | Total | 57 | 2.222 | | | 1 | |
| ***In planta*** | Treatment | 1 | 14.665 | 14.665 | 54.499 | 0.257 | 0.001 |
| | Sampletype | 1 | 0.950 | 0.950 | 3.531 | 0.017 | 0.003 |
| | Water | 1 | 0.352 | 0.352 | 1.307 | 0.006 | 0.165 |
| | Group | 4 | 1.751 | 0.438 | 1.627 | 0.031 | 0.01 |
| | Residuals | 146 | 39.287 | 0.269 | | 0.689 | |
| | Total | 153 | 57.005 | | | 1 | |

[1] degrees of freedom, [2] sum of squares, [3] mean sum of squares, [4] Adjusted p-values are based on 9999 permutations with subsequent.

**Supplemental Table 4.** PERMANOVA analysis of 16S rRNA field trial data. PERMANOVA analysis was conducted using the Bray-Curtis distance of mock and dSynCom samples per sample type, water treatment and inoculation time points.

| **Factor** | **DF**[1] | **SumsOfSqs**[2] | **MeanSqs**[3] | **F Model** | **R2** | **Adjusted p-value**[4] |
|---|---|---|---|---|---|---|
| dSynCom Treatment | 1 | 1.307 | 1.307 | 8.486 | 0.014 | 0.001 |
| Water | 1 | 27.621 | 27.621 | 179.353 | 0.305 | 0.001 |
| Sampletype | 1 | 7.959 | 7.959 | 51.684 | 0.088 | 0.001 |
| Inoculation time | 1 | 0.560 | 0.560 | 3.637 | 0.006 | 0.004 |
| Group | 11 | 6.646 | 0.604 | 3.923 | 0.073 | 0.001 |
| Residuals | 302 | 46.509 | 0.154 | | 0.513 | |
| Total | 317 | 90.603 | | | 1 | |

[1] degrees of freedom, [2] sum of squares, [3] mean sum of squares, [4] Adjusted p-values are based on 9999 permutations with subsequent.

**Supplemental Table 5.** Permutest analysis of 16S rRNA field trial data for each sample type and water treatment by dSynCom inoculation. Permutest analysis was conducted using the Bray-Curtis.

| | | | **CAP analysis, partition of squared Bray distance** | | | **Permutest analysis for all constrained eigenvalues** | | | | |
|---|---|---|---|---|---|---|---|---|---|---|
| **dSynCom inoculation** | **Water treatment** | **Sample type** | | **Inertia[1]** | **Proportion** | | **DF[2]** | **Inertia** | **F Model** | **Adjusted p-value[3]** |
| Onto seeds | Normal irrigation | Root | Constrained | 0.787 | 0.102 | Model | 1 | 0.787 | 4.296 | 0.001 |
| | | | Unconstrained | 6.962 | 0.898 | Residual | 38 | 6.962 | | |
| | | | Total | 7.749 | 1 | | | | | |
| | | Rhizosphere | Constrained | 0.789 | 0.099 | Model | 1 | 0.789 | 4.185 | 0.001 |
| | | | Unconstrained | 7.163 | 0.901 | Residual | 38 | 7.163 | | |
| | | | Total | 7.952 | 1 | | | | | |
| | Drought stress | Root | Constrained | 0.258 | 0.050 | Model | 1 | 0.258 | 2.014 | 0.014 |
| | | | Unconstrained | 4.866 | 0.950 | Residual | 38 | 4.866 | | |
| | | | Total | 5.124 | 1 | | | | | |
| | | Rhizosphere | Constrained | 0.340 | 0.064 | Model | 1 | 0.340 | 2.608 | 0.001 |
| | | | Unconstrained | 4.952 | 0.936 | Residual | 38 | 4.952 | | |
| | | | Total | 5.292 | 1 | | | | | |
| Onto seeds and 4-week old plants | Normal irrigation | Root | Constrained | 0.617 | 0.090 | Model[1] | 1 | 0.617 | 3.755 | 0.001 |
| | | | Unconstrained | 6.242 | 0.910 | Residual | 38 | 6.242 | | |
| | | | Total | 6.859 | 1 | | | | | |
| | | Rhizosphere | Constrained | 0.493 | 0.069 | Model | 1 | 0.493 | 2.688 | 0.001 |
| | | | Unconstrained | 6.597 | 0.931 | Residual | 36 | 6.597 | | |
| | | | Total | 7.090 | 1 | | | | | |
| | Drought stress | Root | Constrained | 0.172 | 0.037 | Model | 1 | 0.172 | 1.411 | 0.096 |
| | | | Unconstrained | 4.514 | 0.963 | Residual | 37 | 4.514 | | |
| | | | Total | 4.686 | 1 | | | | | |
| | | Rhizosphere | Constrained | 0.251 | 0.054 | Model | 1 | 0.251 | 2.173 | 0.005 |
| | | | Unconstrained | 4.382 | 0.946 | Residual | 38 | 4.382 | | |
| | | | Total | 4.632 | 1 | | | | | |

[1] squared Bray distance, [2] degrees of freedom, [3] Adjusted p-values are based on 9999 permutations with subsequent.

**Supplemental Table 6.** PERMANOVA analysis of the root phenotyping data of the field trial. PERMANOVA analysis was conducted using the Euclidean distance of mock and dSynCom samples per water treatment and inoculation time points.

| | **PERMANOVA test** | | | | | | |
|---|---|---|---|---|---|---|---|
| **dSynCom inoculation** | **Factor** | **DF[1]** | **SumsOfSqs[2]** | **MeanSqs[3]** | **F Model** | **R2** | **Adjusted p-value[4]** |
| Onto Seeds | Treatment | 1 | 4.307 | 4.307 | 0.694 | 0.004 | 0.424 |
| | Water | 1 | 635.660 | 635.660 | 102.377 | 0.578 | 0.001 |
| | Group | 1 | 0.873 | 0.873 | 0.141 | 0.001 | 0.849 |
| | Residuals | 74 | 459.468 | 6.209 | | 0.418 | |
| | Total | 77 | 1100.309 | | | 1 | |
| Onto seeds and 4-weeks old plants | Treatment | 1 | 8.061 | 8.061 | 1.691 | 0.010 | 0.202 |
| | Water | 1 | 444.685 | 444.685 | 93.301 | 0.540 | 0.001 |
| | Group | 1 | 9.279 | 9.279 | 1.947 | 0.011 | 0.166 |
| | Residuals | 76 | 362.226 | 4.766 | | 0.439 | |
| | Total | 79 | 824.251 | | | 1 | |

[1] degrees of freedom, [2] sum of squares, [3] mean sum of squares, [4] Adjusted p-values are based on 9999 permutations with subsequent.

**Supplemental Table 7.** Pairwise PERMANOVA analysis of the root phenotyping data of the field trial.

| | Pairwise PERMANOVA | | | |
|---|---|---|---|---|
| **dSynCom inoculation** | **Factor 1** | **Factor 2** | **R2** | **Adjusted p-value[1]** |
| Onto seeds | Mock Drought | SynCom Drought | 0.006 | 5.322 |
| | Mock Watered | SynCom Watered | 0.021 | 2.652 |
| Onto seeds and 4-weeks old plants | Mock Drought | SynCom Drought | 0.027 | 2.088 |
| | Mock Watered | SynCom Watered | 0.113 | 0.036 |

[1] Adjusted p-values are based on 9999 permutations with subsequent.
